# Evidence for PKD2L1-positive neurons present distant from the central canal in the ventromedial spinal cord and *Medulla* of the adult mouse

**DOI:** 10.1101/2021.05.18.440679

**Authors:** Nina Jurčić, Caroline Michelle, Jérôme Trouslard, Nicolas Wanaverbecq, Anne Kastner

## Abstract

Neurons in contact with the cerebrospinal fluid (CSF) are found around the medullo-spinal central canal (CC) in adult mice. These neurons (CSF-cNs), located within or below the ependymal cell layer known as the stem cell niche, present a characteristic morphology with a dendrite projecting to the CC and ending with a protrusion. They are GABAergic, characterized by an immature neuronal phenotype and selectively express PKD2L1, a channel member of the TRP channel superfamily with properties of sensory receptor.

Using immunohistological techniques in mice, we characterize a new population of PKD2L1 positive cells that is observed around embryonic day 16 (E16), is present distant from the CC in a zone enriched with astrocytes and ependymal fibers of the ventro-medial spinal cord and medulla. With development, their number appears stable although smaller than that of CSF-cNs and they progressively become more distant from the CC with the reorganization of the CC region. These neurons share both functional and phenotypical properties with CSF-cNs, but they appear subdivided in two groups. One, present along the midline, has a bipolar morphology and extend a long dendrite along ependymal fibers and towards the CC. The second group, localized in more ventro-lateral regions, has a multipolar morphology and no apparent projection to the CC

Altogether, we describe a novel population of PKD2L1^+^ neurons distant from the CC but with properties similar to CSF-cNs that might serve to sense modification in the composition of either CSF or interstitial liquid, a function that will need to be confirmed.

## INTRODUCTION

The central canal (CC) in the spinal cord (SC) and Medulla (Med) is surrounded by a layer of ependymal cells among which neural stem cells, located mainly at the dorsal and ventral poles (Hugnot and Franzen, 2011; Becker et al., 2018). These latter exhibit long radial processes projecting along the midline and are characterized by the expression of neural stem cells markers as nestin, 3CB2, FoxJ1, Sox2 (Meletis et al., 2008; Hamilton et al., 2009; Marichal et al., 2012). These ependymal stem cells are quiescent in adult mammals (Sabourin et al., 2009) but their proliferation was shown to be triggered following SC injury (Meletis et al., 2008). In addition to stem cells, the ependymal zone (EZ) also contains specific neurons in contact with the cerebrospinal fluid (CSF) (Agduhr, 1922; Bruni and Reddy, 1987; Bjugn et al., 1988; Vigh and Vigh-Teichmann, 1998; Vigh et al., 2004). CSF-contacting neurons (CSF-cNs) have been shown to contact the CSF through a short neurite (10-40 μm length) that crosses the ependymal cell layer and exhibits a terminal bud within the lumen (Orts-Del’immagine et al., 2012; Orts-Del’Immagine et al., 2014). Their axonal processes appear to project medially toward the ventral fissure of the SC where they regroups in longitudinal fiber bundles (Stoeckel et al., 2003). One striking feature for CSF-cNs, conserved in all vertebrates, is the selective expression of the PKD2L1 (polycystic kidney disease 2-like 1), a cationic non-selective channel belonging to the TRP superfamily that contributes to their sensory function. In particular PKD2L1 is capable of detecting and integrating changes in extracellular pH and osmolarity (Huang et al., 2006; Orts-Del’immagine et al., 2012; Orts-Del’Immagine et al., 2014; Jalalvand et al., 2016a, 2016b) as well as regulating spinal motor network activity and locomotion (Böhm et al., 2016; Sternberg et al., 2018). Based on the studies conducted in lower vertebrates, CSF-cNs were proposed to represent a novel sensory system intrinsic to the central nervous system (CNS) and capable of modulating its activity. On the other hand, the CSF was shown to play an important role during CNS development by releasing bioactive molecules (growth factors and morphogens) thus promoting neurogenesis, neuronal differentiation and migration. Moreover, following pathological states such as neuroinflammation and lesion, CSF composition is modified and enriched with inflammatory factors (Kothur et al., 2016; Simon and Iliff, 2016; Bjorefeldt et al., 2018). Altogether, when considering the strategical position of CSF-cNs between CSF and parenchyma and the role for CSF, one can suggest that this unique neuronal population might play an important role in informing about CNS state and pathological dysfunction. Finally, although CSF-cN neurogenesis has been shown to occur only during development they have been shown to retain an incomplete maturity state (Stoeckel et al., 2003; Sabourin et al., 2009; Orts-Del’Immagine et al., 2014, 2017) and might be reactivated and take part to plasticity and regenerative processes associated with the ependymal stem cell niche.

Most studies focused on ependymal and sub-ependymal PKD2L1^+^ CSF-cNs (located at less than 50 μm from the CC) however previous observations (Orts-Del’immagine et al., 2012; Djenoune et al., 2014; Orts-Del’Immagine et al., 2014), recently confirmed (Tonelli Gombalová et al., 2020), mentioned the presence of PKD2L1^+^ cells located more distally from the CC. The developmental origin for these distal PKD2L1^+^ cells is still unclear but as ependymal cells located along the ventral midline in regions distal from the CC display specific electrophysiological properties of neuronal progenitor (eg. Ca^2+^ spikes), they have been suggested to represent potential CSF-cN precursors (Marichal et al., 2012). This assumption is further supported by a recent study from DiBella and colleagues (2019) showing that expression of Ascl1, the basic helix-loop-helix (bHLH) transcription factor, triggers CSF-cN production from ependymal cells. Nevertheless, until recently (Tonelli Gombalová et al., 2020), distant PKD2L1 cells have never been studied and little is known about their origin, properties and interaction with ependymal cells. Thus, a better characterization of these distal PKD2L1 expressing cells appears crucial to demonstrate their neuronal nature, their properties and whether they are in contact with the CSF as well as integrated to the CSF-cN network.

In the present report, we describe the morphology, distribution and organization of distal PKD2L1 expressing cells in the spinal cord (SC) and *Medulla* (Med). We indicate that they are neurons, share features with CSF-cNs and we characterize their relationship with the CC and the astroglial-ependymal stem cell niche. This study provides new insights to better understand the organization of PKD2L1^+^ neurons around the CC and in in the ventro-medial zone of the spinal cord and medulla and sets ground to investigate in the future their interaction with the ependymal stem cell niche and their function.

## MATERIAL & METHODS

### Animals and ethical statement

All experiments were conducted in conformity with the rules set by the EC Council Directive (2010/63/UE) and the French ‘Direction Départementale de la Protection des Populations des Bouches-du-Rhône’ (Project License Nr: APAFIS 17596; 2018111919329153. N.W. and License for the use of Transgenic Animal Models Nr: DUO-5214). Protocols used are in agreement with the rules set by the Comité d’Ethique de Marseille, our local Committee for Animal Care and Research. All efforts were made to ensure animal well-being and minimize animal suffering and the number of animals used. Animals were housed at constant temperature (21°C) under a standard 12 h light-12 h dark cycle, with food (pellet AO4, UAR, Villemoisson-sur-Orge, France) and water provided ad libitum. Experiments were carried out on adult (3 months old: 3M, n = 8), neonate (postnatal days 0-2: P0-P2, n = 6) and embryonic (E15, 16 and 18, n = 3 to 6) male and female wild-type (C57BL/6J) and transgenic mice. We took advantage of the PKD2L1-IRES-Cre mouse model (generous gift from Dr CS Zuker) to generate through cross-breeding mice selectively expressing in PKD2L1 cells EGFP (Z/EG reporter transgenic mice provided by Dr P Durbec; PKD-Cre::flex-EGFP, PKD2L1-GFP) or tdTomato (Gt(ROSA)26Sor^tm14(CAG-tdTomato)Hze^ from the Jackson laboratory; PKD-Cre::flex-tdTomato, PKD-tdTomato) cytosolic fluorescent markers. Thus, in PKD2L1^+^ cells, EGFP or tdTomato expression was selectively induced by Cre recombinase activity (Huang et al., 2006). To assess EGFP and tdTomato expression in PKD2L1^+^ animals, we carried out PCR on tail genomic DNA as specified in our previous work (Orts-Del’immagine et al., 2012).

### Immunohistochemistry and antibody specificity

Prior to the procedure (30 min), adult mice were injected with Metacam (5 m.Kg^−1^), a non-steroidian anti-inflammatory compound. Subsequently, they were anaesthetized with intraperitoneal (IP) injection of ketamine - xylazine mixture (100 and 15 mg/kg, respectively) and injected on the site of section with Lurocaine (5 m.Kg^−1^), a local antalgic. Next, animals were transcardially perfused first with phosphate buffer solution (PBS 0.1M) and subsequently with 4% paraformaldehyde (PFA) in PBS (PBS–4% PFA). For the experiments where GABA immunoreactivity was tested, mice were transcardially perfused with a fixation medium containing 4% PFA and 0.2% glutaraldehyde. Brain and spinal cord (SC) tissues were immediately removed and post-fixed 1 h in PBS–4% PFA at 4°C. Postnatal mice (P0-2; wild type and PKD-tdTomato) were anaesthetized using progressive cooling to 0°C, the medullo-spinal tissue dissected and immersed in PBS–4% PFA for 18-24h at 4°C. Mice embryos at embryonic days 15 to 18 (E15-18) were isolated from 6 to 8-week-old time-pregnant PKD-tdTomato female. Briefly, following anesthesia (see above), the mother was sacrificed and its abdomen cut-opened after injection of local antalgic solution (Lurocaine). The isolated embryos were then immersed in in PBS–4% PFA for 18-24h at 4°C. All tissues were subsequently cryoprotected for 24–48 h in 30% sucrose at 4°C and frozen in isopentane (−40°C). For adult animals, we distinguished two regions for the brainstem or *Medulla* (Med): caudal (cMed, from −8.20 to −7.90 mm) and sub-postremal (spMed, from −7.90 to −7.65 mm) *Medulla* as well as two regions for the spinal cord (SC): cervical C1 SC region (cervSC, antero-posterior stereotaxic coordinates of regions more caudal than −8.50 mm from Bregma; Paxinos Mouse Atlas) and lumbar SC (lumbSC). For P0 we only considered and compared the regions of the Med and of the cervSC while at embryonic stages only the rostral SC. Coronal and sagittal thin sections (30 μm) were obtained using a cryostat (Leika CM3050) and collected serially in 12-well plates containing 0.1M PBS or directly on slides for embryo and p0 mice. Sections were incubated first 1h in PBS containing 0.3% Triton X-100 and then in a blocking solution with 3% horse or goat normal serum and 1% bovine serum albumin (BSA). Sections were incubated overnight at 4°C with primary antibodies and 3% horse or goat normal serum. Sections were then washed in PBS and incubated for 2h with secondary antibodies conjugated to either AlexaFluor 488 or 594 (1:400, Life Technologies) and then washed again. In double immunolabelling experiments, the two primary antibodies were generally applied sequentially. Sections were mounted on gelatine coated slides and coverslipped with Mowiol mounting medium for fluorescence microscope preparation. In some instance, nuclear labelling was also performed using DAPI staining (1/1000, Sigma D9542). To assess the selectivity of the observed immunolabeling, in each set of experiments, some sections were treated in the absence of primary antibodies (negative control). Typically, each experiment was performed on tissues obtained from either 2-3 different 3-month-old animals, 3 postnatal mice (P0-2) or 3 to 6 embryos at each embryonic age. The primary antibodies used for this study are: DCX (Santa Cruz SC-8066, goat, 1:100), GABA (Immunostar 20094, rabbit, 1:250), GAD67 (Millipore MAB5406, rabbit, 1:500), GFAP (Sigma, G3893, mouse, 1:500) GFP (Gift from Dr T Doan, rabbit, 1:7500), HuC/D (Molecular Probes A-21271, mouse, 1:1000), NF160 (Sigma, N5264, mouse, 1:80), MAP2 (Sigma M-1406, mouse, 1:600), NeuN (Millipore MAB377, mouse, 1:800), Nkx6.1 (Dev.Stud. Hybrid.Bank F55A10, mouse, 1:100, gift of Dr JP Hugnot), PSA-NCAM (Millipore MAB5324, mouse IgM, 1:250) PKD2L1 (Millipore AB9084, rabbit, 1:700) and Vimentin (Millipore AB5733, chicken, 1/500).

All antibodies have been used in our previous works (Orts-Del’Immagine et al., 2014, 2017). For NeuN, MAP2, HuC/D, the antibody specificity was assessed by the observation of the expected labelling in neuronal cells visualized in the same section. Vimentin and GFAP immunofluorescence were detected as expected in ependymal cells and astrocytes. The specificity of PKD2L1 immunolabelling was confirmed by the use of the PKD2L1-GFP and PKD-tdTomato mouse lines. Concerning Nkx6.1, the selected antibody is currently used for the detection of this transcription factor in neuronal progenitors and was successfully used by other authors. DCX and PSA-NCAM antibody specificity was confirmed in previous reports by their selective labelling of neuroblasts in the SVZ and Dentate gyrus (Orts-Del’Immagine et al., 2017). Moreover, the presence of DCX in CSF-cNs was confirmed by the use of a DCX-GFP mice (Orts-Del’Immagine et al., 2017). The selectivity of the GABA/GAD67 immunolabeling was confirmed by the presence of GABA/GAD67 cells in brain regions known to contain GABAergic neurons: the spinal cord dorsal horn, the *nucleus tractus solitari* and the cerebellum (Orts-Del’Immagine et al., 2017).

### Image Acquisition and analysis

Sections were observed on a confocal laser scanning microscope (CLSM; Zeiss LSM700) equipped with solid state fiber optic lasers and single plane images or stacks of images were acquired with a 2x digital zoom using a 20x objective (numerical aperture, NA: 0.8 for an optical thickness of ∼2 μm). Images were acquired at optimal resolution between 512×512 and 1024×1024 pixels. When using more than one fluorochromes with different excitation/emission spectra, images were acquired sequentially for each channel (405 nm, blue, 488 nm, green and 555 nm, red) and the filters and photomultiplier tubes settings chosen to optimize images and avoid signal crosstalk. Images were analyzed and prepared using ZEN 2009 light Edition (Zeiss software), ImageJ 1.45 (NIH) and Corel Photo-Paint. For better visualization, contrast and brightness were adjusted in images used for the figures. For analysis of cell density, PKD2L1^+^ cells were counted manually in each image and the total cell number divided by the section thickness (10 to 20 μm stack depth). For the analysis of CC parameters, PKD2L1^+^ cell distribution, distance to the CC and to the ventral midline (expressed in μm) measurements were carried out using ImageJ. At each medullo-spinal level, the analysis was performed in at least 2 animals and a minimum of 6 different sections per animal. To analyze the intensity of PKD2L1 immunoreactivity (IR) in each cell as a function of their distance from the CC, IR intensity (LUT on grey scale) was measured in a small region on the soma (10-15 μm^2^) cell by cell on each section (in the cervSC of 3 mice) and subtracted from the background intensity. Typically, analyzed images had 8 bits grey scale resolution (values from 0 to 256) and fluorescence intensity was comprised between 0 bit (black) and 256 bits (white and saturated), the average grey scale value of the background intensity being of 220. For each images, the number of optical sections used for the Z-projection is indicated in the figure legends.

### Slice preparation and electrophysiology

Coronal brainstem slices were prepared from 3-months-old PKD2L1:EGFP mice as previously described by (Orts-Del’immagine et al., 2012). Briefly, the animal was anesthetized with an IP injection of a ketamine and xylazine mixture (see above, Immunohistochemistry). Following decapitation, brain was removed and immersed in ice-cold (0-4 °C), oxygenated (95% O_2_, 5% CO_2_), low calcium/high magnesium slicing artificial cerebrospinal fluid (S-aCSF) solution containing (in mM): NaCl 75, NaH_2_PO_4_ 1.25, NaHCO_3_ 33, KCl 3, MgCl_2_ 7, sucrose 58, glucose 15, ascorbic acid 2, myo-inositol 3, sodium pyruvate 2, CaCl_2_ 0.5 (pH 7.35-7.40, osmolality of 300-310 mosmol.kg^−1^). Coronal brainstem slices (250 μm thick) were cut using a vibratome (Leica VT1000S) and incubated at 35 °C for 15-20 min in a submerged recovery chamber filled with oxygenated recording aCSF (R-aCSF) containing (in mM): NaCl 115, NaH_2_PO_4_ 1.25, NaHCO_3_ 26, KCl 3, MgCl_2_ 2, glucose 15, ascorbic acid 2, myo-inositol 3, sodium pyruvate 2, CaCl_2_ 2 (pH 7.35-7.40, osmolality of 300-310 mosmol.kg-1). Slices were left to recover for 1h from 35 °C to room temperature under continuous oxygenation. After a recovery period, slices were transferred one by one to the recoding chamber superfused with R-aCSF at 1.5-2.5 ml min^−1^. Slices in the recording chamber were visualized under infrared DIC (IR-DIC) illumination mounted on a Zeiss Axioscope S1 microscope. GFP^+^ cells in ventro-medial Medulla were visualized with a 480 nm excitation light using p1 precisExcite LED epifluorescence system (CoolLED, Roper) and a CoolSNAP HQ2 cooled CCD camera (Photometrics) connected to a computer through a frame grabber (CoolSNAP LVDS interface cards, Photometrics) and controlled by MetaView software (Molecular Devices Inc). Whole-cell patch-clamp recordings were performed at room temperature in voltage- and current-clamp mode using Multiclamp 700B patch-clamp amplifier connected to a Digidata 1322A interface (Molecular Devices Inc). Patch pipettes (4-6 MΩ) were pulled from borosilicate glass capillaries (Harvard Apparatus) using a horizontal P-97 Flaming/Brown type micropipette puller (Sutter Instrument Co). Patch pipette was filled with the intracellular solution containing (in mM): potassium gluconate 120, NaCl 5, HEPES 10, MgCl_2_ 1, CaCl_2_ 0.25, EGTA 2, Mg-ATP 4, disodium phosphocreatine 10, Na_3_GTP 0.2 (adjusted to pH 7.33 with KOH; osmolality of 295 mosmol.kg^−1^). AlexaFluor 594 (10 μM, excitation 590 nm/emission 610 nm, Invitrogen) was added to the intracellular solution in all the experiments to visualize the morphology of patched neuron (Orts-Del’immagine et al., 2012). In the whole-cell configuration, series resistance (r*s*) was between 10-20 MΩ and was monitored throughout each experiment by applying a −20 mV calibration pulse.

Recordings were discontinued when r*s* exceeded by more that 25% the original value. Signals were filtered at 2.4 kHz, digitized at 10 kHz and acquired on a computer using the Clampex 9.2 software (Molecular Devices Inc.). Single-channel and synaptic activities were recorded in voltage-clamp mode from a holding potential of −70 mV. Current-clamp recordings were performed from the resting membrane potential (RMP) and action potentials (APs) triggered with 500 ms direct current (DC) pulses from −10 pA to +30 pA, with 10 pA increments in the presence of oxygenated R-aCSF supplemented with 10 μM gabazine and 1 μM strychnine (GABA_A_ and glycine receptor antagonists, respectively; Sigma-Aldrich).

Current responses were analyzed using Clampfit 10.0 suite (Molecular Devices Inc.) and Excel 2016 (Microsoft). RMP was determined in current-clamp mode at I = 0 just after the whole-cell configuration was achieved as well as by averaging in the current-clamp recordings a ~1 s period in the absence of current injection (injection = 0 pA). AP properties were analyzed from the recordings of spontaneous discharge activity (10-30 s) or from trains of APs elicited by +30 pA current injections for 500 ms using the “Threshold Search” routine from Clampfit 10.0 with the threshold set at 0 mV. For each condition, principal parameters of APs (amplitude, width, frequency) were determined. AP threshold potential (TP) was determined by averaging the voltage response during current (I injection, +30 pA) and before the AP raising phase. Single-channel activity was analyzed from spontaneous current traces (120 s) obtained in voltage-clamp mode using the “Single-Channel Search” routine from Clampfit 10.0. For each recording, the unitary current amplitude, channel open probability, and frequency were determined. Channel open probability (N.Po) was measured as the sum of open times divided by the time of analysis for each condition. The frequency of channel opening was calculated as the number of detected events divided by the time of analysis. Data are expressed as means ± SD.

### Statistical analysis and Data representation

All data are expressed as mean ± SD and represented as with whisker plots using the Tukey’s method as well as ‘violin’ plots to illustrate data point density. In whisker plots, for each data set, the median and the 25th to the 75th percentiles (lower and upper limits of each bar, respectively) are calculated. Next, the interquartile distance (IQR) is determined as the difference between the 25th and 75th percentiles and the whiskers limits or ‘inner fences’ calculated as the 75th percentile plus 1.5 times IQR and the 25th percentile minus 1.5 times IQR. All data with value either higher or lower than the inner fences are represented as individual data points and considered as outliers (circles on the plots). The analyses of the cell number, cell types around the CC (CSF-cNs) and ventral from it (dPKD2L1 neurons), cell distance to the CC or from midline and phenotypical characteristics (NeuN and DCX percentage of expression) were collected in adult animals as a function of 4 medullo-spinal regions (lumb and cervSC, c and spMed; Region). For the analysis and comparison between ages at P0-2 and 3M (Age), we only distinguished *Medulla* (cMed) and cervical (cervSC) in 2 to 3 animals. In the comparison of the various parameters during development between E15 and 3M, we analyzed cervical tissues. The distinction between CSF-cNs and distal PKD2L1 neurons is considered the cell type (Type) variable while the distinction between the different levels of the CC rostro-caudal axis represents the Region variable. On one hand, in a given animal of a given age, the organization of each Region and neurons’ Type is assumed dependent from the organization of the other ones and the variable represents dependent factors. On the other hand, the same variables represent independent factors when considered as a function of Age and in different animals. Therefore, the statistical analyses required using a mixed effect model taking into account in a hierarchical way both dependent (Region, Type) and independent variables (Age) as well as the interaction between the variables. Finally, the choice of animals used for this study was done randomly among all those available, therefore it is necessary to implement the statistical model with a factor corresponding to this random effect. The statistical analyses were carried out using the R Studio 3.3.1 statistical software (R Studio Team, 2015) in the R environment and the “*non-linear mixed effect* (nlme version 3.1-128)” package (Pinheiro et al., 2016). The null hypothesis was set as the absence of difference between the data among each group. The *nlme* model was tested by using an analysis of the variance (ANOVA) and a subsequent post-hoc Sidak test for multiple pairwise comparison was used. We provide the values for χ^2^, the degree of freedom (number of parameter −1,dF) and of the p-values for given fixed factor tested (Region, Type or Age). Wherever, an interaction between the Region, Type or Region and Age factors was statistically significant we reported the p-value. Statistical differences were considered as significant for p < 0.05. For results where data are only compared for one variable, we used one-way ANOVA statistical analysis and provide F-statistic, degree of freedom, R^2^ and p-value for the analyzed factor.

## RESULTS

### PKD2L1 expressing cells are present in the ventro-medial spinal cord and Medulla

In previous studies, we have described, in adult mice, and characterized PKD2L1^+^ neurons around the CC in the cervSC and Med (Orts-Del’immagine et al., 2012, 2014, 2016 and 2017) that correspond to CSF-cNs as described in several vertebrate models (Agduhr, 1922; Vigh and Vigh-Teichmann, 1998; Vigh et al., 2004; Djenoune et al., 2014; Jalalvand et al., 2016a). Here, using immunostaining against PKD2L1 on transverse and sagittal cervSC sections, we show that PKD2L1 positive cells can also be found outside the peri-ependymal zone (EZ) even at long distance from the CC (Fig. 1A, B), principally in the ventromedial region. In the rest of the study, we will distinguish PKD2L1^+^ cells present around the CC, designed as CSF-cNs, from those sitting at a distance greater that 50 μm named distal(d) cells (dPKD2L1 cells). Figure 1B further illustrates on a cervSC sagittal section that dPKD2L1 cells spread across the whole grey matter, are localized as ventral as in the white matter (asterisk in Fig. 1B) and that long PKD2L1^+^ fibers projecting in the dorso-ventral axis can be observed, presumably corresponding to dPKD2L1 dendritic processes (Fig. 1A and B, black arrows and see below). Although dPKD2L1 cells are observed as single cells mainly bipolar, scattered in the parenchyma, we observe also observed, along the rostro-caudal axis, dPKD2L1 cells with a more complex morphology cells and organized in patches/clusters regularly spaced (Fig. 1B, white arrows, see Fig. 2 and below). Looking at the somatic PKD2L1 IR, we noticed that it appears to decrease with the increase in the distance of dPKD2L1 cells from the CC. Thus, in the cervSC, the average PKD2L1 IR intensity was 115 ± 4.1 a.u. for cells at less than 200 μm from the CC and around half (70.5 ± 4.5 a.u.) for cells further away (p = 9.3.10^−12^) suggesting that dPKD2L1 in more ventral region of the CC would have a lower PKD2L1 channel expression. Next, we analyzed the distribution of dPKD2L1 cells along the CC axis at the lumbar level of the SC (lumbSC, Fig 1C) as well as the caudal (cMed, Fig 1E) and subpostremal (spMed, Fig 1F) levels in the Med. Z-projections of ~10 serial transverse sections indicate that dPKD2L1 cells are present at all levels where they mainly reside ventro-medially to the CC, even in the white matter within a sector covering approximately 400 μm x 300 μm (Fig. 1A and C-F). Note that distal cells are also found in thoracic SC (data not shown). We then determined the number of dPKD2L1 cells present in the medullo-spinal tissue compared to CSF-cNs and, on average, we found 3.5 ± 0.4 dPKD2L1 cells and 9.2 ± 0.2 CSF-cNs across 10 μm of tissue depth (Fig. 1G; 3 animals and 16, 33, 33 and 19 sections for lumbSC, cervSC, cM and spM, respectively; *nlme* model: χ^2^(Type) = 289.55, dF = 1, p(χ^2^) < 2.10^−16^). The analysis of dPKD2L1 cell density within each of the 4 regions of interest indicates that, at all levels, dPKD2L1 cell number is low but homogenous across regions (Fig. 1G; lumbSC: 3.9 ± 1.2; cervSC: 3.6 ± 1.5; cMed: 3.3 ± 1.5 and spMed: 3.1 ± 1.6 cells/10 μm tissue depth, 3 animals and 16, 33, 33 and 19 sections; *nlme* model: χ^2^(Region) = 11.12, dF = 3, p(χ^2^) = 0.011). When compared to the distribution of CSF-cNs, dPKD2L1 cell density was smaller in all regions (Fig. 1G; CSF-cNs: lumbSC: 8.8 ± 0.7; cervSC: 10.3 ± 3.6; cMed: 9.7 ± 0.5 and spMed: 7.6 ± 0.7 cells/10 μm tissue depth, 3 animals and 16, 33, 33 and 19 sections) and represented on average 28 ± 2% of the total PKD2L1^+^ cell population (lumbSC: 30.9 ± 7.8%; cervSC: 27.1 ± 8.5%; cMed: 25.7 ± 7.9% and spMed: 28.3 ± 7.6%; 3 animals; One-Way ANOVA: F(Type) = 1.6206, dF = 3, p(F) = 1.1896). We next determined dPKD2L1 cell distribution as a function of the distance from the CC in the dorso-ventral axis and found that the average distance was 163 ± 96 μm with cells present at distances as far as ~500 μm. When analyzing the distance to the CC as a function of the 4 different regions, we noticed that this parameter increased in the caudo-rostral axis and was the highest in the spMed (Fig 1H; 95 ± 34 μm, 168 ± 92 μm, 166 ± 85 and 239 ± 108 μm in lumbSC (n = 69 cells), cervSC (n = 167), cMed (n = 128) and spMed (n = 105), respectively; 3 animals and 16, 33, 33 and 19 sections, respectively; One-Way ANOVA: F(Region) = 37.891, dF = 3, p(F) < 2.2.10^−16^). At the lumbSC level, 81% of dPKD2L1 cells were found in the first 125 μm ventral from the CC and only 34%, 34% and 14% at the cervSc, cMed and spMed, respectively. In contrast, the number of dPKD2L1 cells found at distances higher than 200 μm increases in the caudo-rostral axis with only 1% at lumbSC, 34% at cervSC, 25% at cMed and 60% at spMed. The length of the region ventral to the CC increases from the lumbSC to the spMed (lumbSC: 779 ± 80 μm; cervSC: 835 ± 92 μm; cMed: 844 ± 50 μm and spMed: 1129 ± 113 μm; 3 animals and 16, 33, 33 and 19 sections, respectively). We calculated the ratio between dPKD2L1 position to the CC over the length of the ventral pole, in order to determine whether these two parameters were correlated. Our data indicate that this ratio increases between the lumbSC and the cervSC and then remains stable for more rostral region (lumbSC: 0.12 ± 0.04 μm; cervSC: 0.20 ± 0.11 μm; cMed: 0.20 ± 0.10 μm and spMed: 0.21 ± 0.10; 3 animals, 16, 33, 33 and 19 sections and 133, 167, 128 and 105 cells, respectively; One-Way ANOVA: F(Region) = 25.159, dF = 3, p(F) = 3.144.10^−15^). This result suggests that the increased distance from the CC observed for dPKD2L1 neurons is not due to an enlarged ventral zone of the medullo-spinal tissue. Finally, we show that on average dPKD2L1 cells were localized at 43 ± 53 μm (n = 510 cells) lateral to the ventro-medial axis (midline) and our results indicate that 38% of the cells are distributed at less than 15 μm lateral to the midline while 72% were found at a lateral position within the 50 μm range. On average, dPKD2L1 cells were localized at 14 ± 16 μm, 43 ± 54 μm, 52 ± 48 μm and 70 ± 69 μm in lumbSC (n = 127 cells), cervSC (n = 158 cells), cMed (n = 128 cells) and spMed (n = 97 cells), regions, respectively (Fig 1I; 2-3 animals; One-Way ANOVA: F(Region) = 25.075, dF = 3, p(F) = 3.867.10^−15^). Again, at the lumbSC level the majority of dPKD2L1 cells were found close to the midline in the first 50 μm (lumbSC: 93%, cervSC: 73%, cMed: 61% and spMed: 52%). Thus, along the caudo-rostral axis the dPKD2L1 cell distance from the midline appears to increase.

**Fig. 1:**
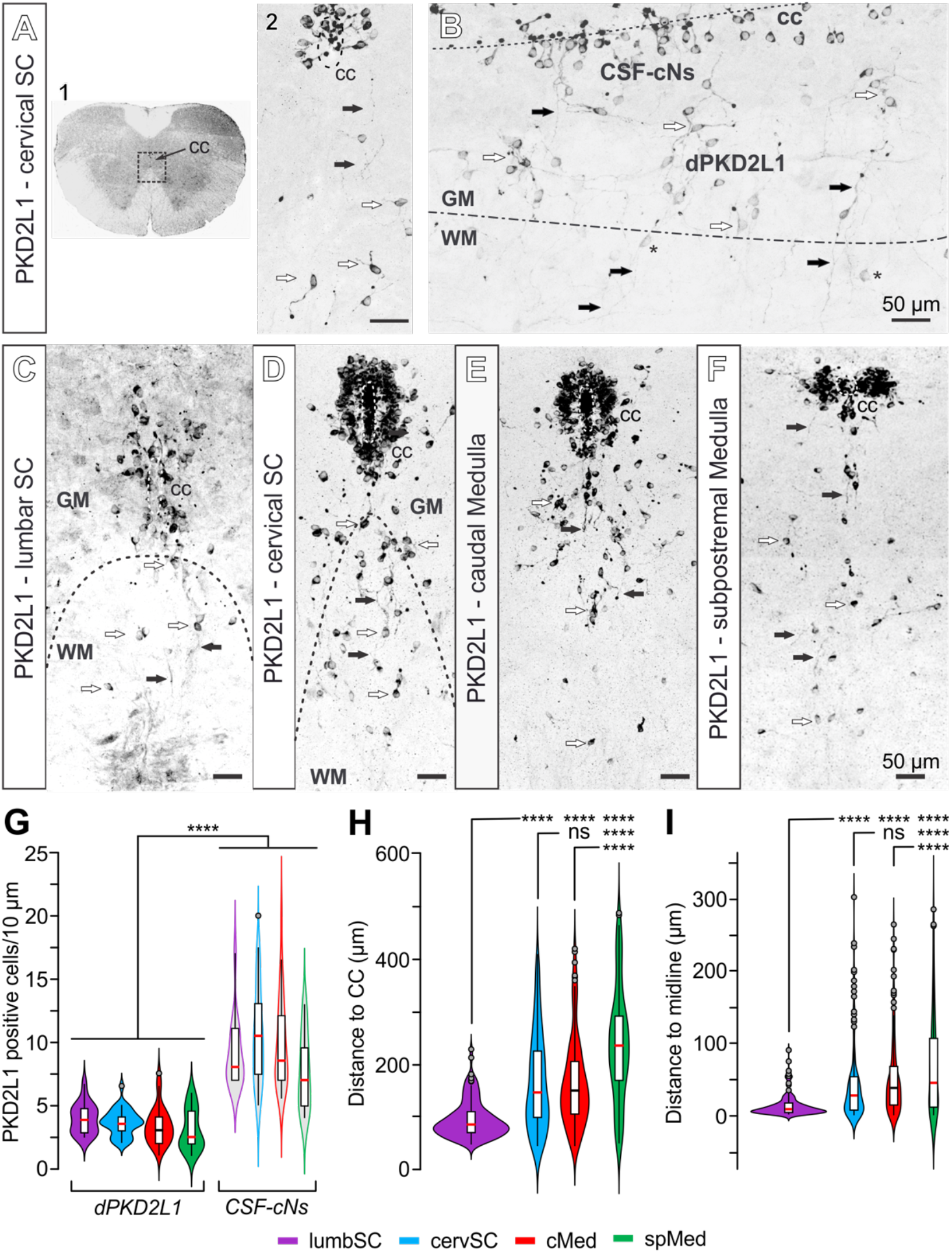
Distal PKD2L1^+^ cells are present in the ventro-medial spinal cord and Medulla. **A1.** PKD2L1 immunolabelling of coronal sections of the cervSC indicating the region (square) where PKD2L1^+^ cells are detected. **A2.** At higher magnification, PKD2L1^+^ cells are found around the CC (CSF-cNs) but also cells present at higher distances (dPKD2L1 cells) in the ventro-medial region **B.** Representative cervSC sagittal section showing dPKD2L1 cell organization in the ventro-medial region (PKD2L1 immunolabelling). **C-F.** Projection from serial PKD2L1 immunolabelled coronal sections at the 4 levels of interest showing the distribution of PKD2L1^+^ cells around the CC and in the ventral region: lumbar SC (C, 5 sections), cervSC, caudal and subpostremal Medulla (D-F, 10 serial sections). **G.** Summary whiskers box plots comparing the numbers of dPKD2L1 cells (filled colored bars, left) and CSF-cNs (colored outlines, right) in the 4 analyzed regions. **H** and **I.** Summary whiskers box plots of dPKD2L1 cell distance from the CC (H) and from the midline (I) for the 4 regions of interest. The data point density is shown with violin plots and the same color code for the regions of analysis (lumbSC, cervSC, cMed and spMed) is used for G, H and I. Dotted-Dashed lines: limit between the grey (GM) and white matter (WM); Dashed lines: CC border; White arrows: dPKD2L1^+^ cells; Black arrows: dPKD2L1^+^ processes. Scale bar: 50 μm (A1, B-F); Z-projections of 5 (A2) and 15 (B) optical sections.

**Fig. 2:**
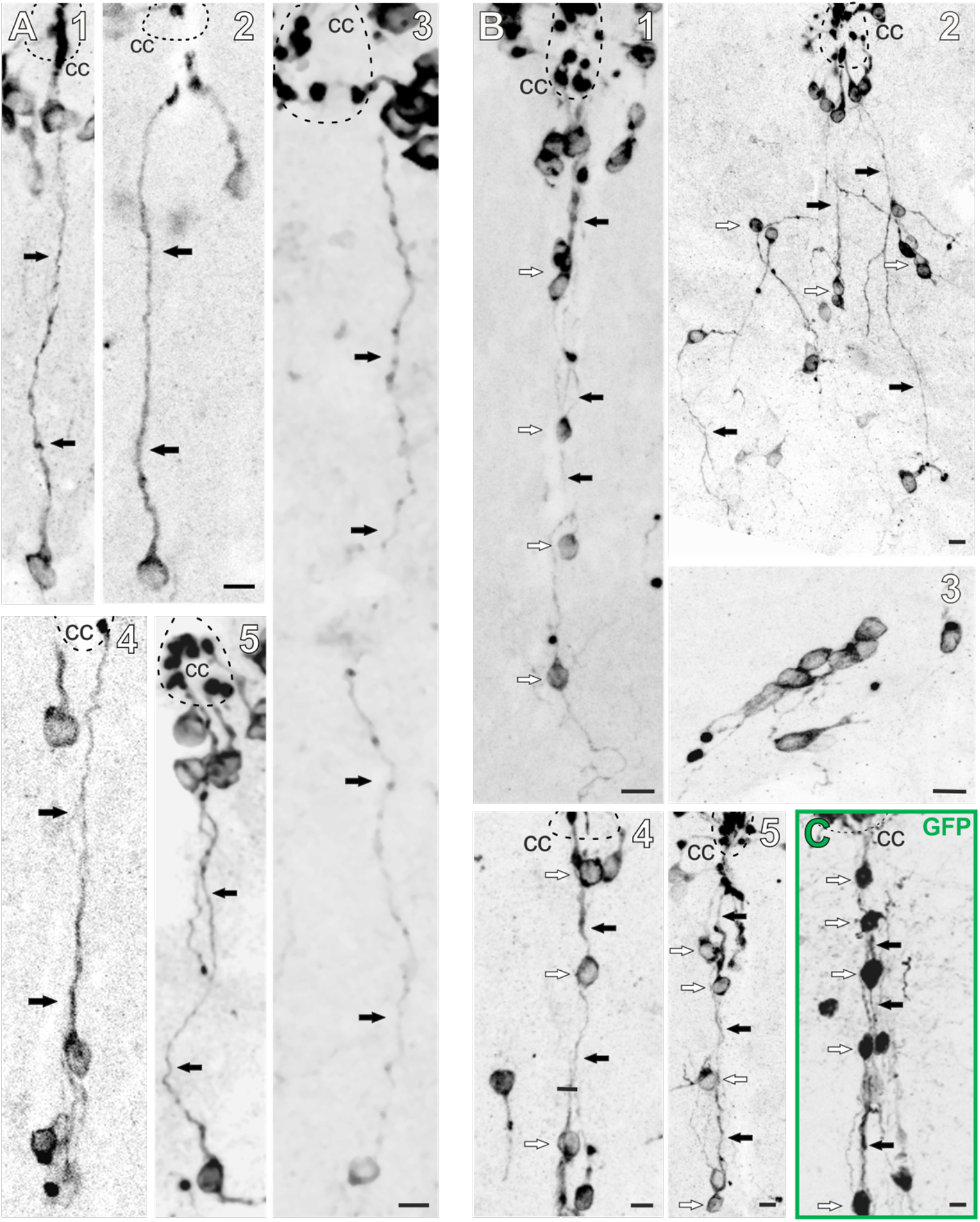
dPKD2L1 cells project to the CC with long projections or form cell chains. **A.** Examples of PKD2L1 immunolabelled coronal cervSC sections showing dPKD2L1 cells with long dorso-ventral processes (80 to 400 μm) reaching the CC (A1-A5). For A2-4 note the presence of a typical bud in contact with the CC. **B.** Examples of PKD2L1 immunolabelled cervSC sections showing typical cell chain/cluster organization of dPKD2L1 cells. **B1, 4 and 5**. Examples of dPKD2L1 cells forming long ventro-medial cell chains towards the CC in the cervSC level. **B2.** Example of a section (cervSC level) illustrating bipolar dPKD2L1cells with long processes directed toward the CC and PKD2L1^+^ cells with a multipolar morphology. **B3.** Micrograph illustrating the typical cluster formed by dPKD2L1 cells. **C.** Example of a GFP immunolabelled section (Medulla) from a PKD2L1::GFP mouse (in which GFP reveals the whole processes of PKD2L1 expressing cells) showing the dPKD2L1 cell chain organization. Dashed lines: CC borders; White arrows: dPKD2L1^+^ cells; Black arrows: dPKD2L1^+^ processes. Scale bar: 10 μm (A-C); Z-projections of 3-5 (A) and 13, 15 (B, C) optical sections.

### Are dPKD2L1 in contact with the central canal?

Around the CC, CSF-cNs have been shown to have a characteristic morphology with a small soma (diameter ~10 μm) sitting within or close to the ependymal cell layer. They project a short dendrite (10-40 μm length, MAP2 positive) that crosses the ependymal cell layer and ends in the CC lumen with a terminal protrusion (bud) exhibiting a strong PKD2L1 IR (Agduhr, 1922; Vigh and Vigh-Teichmann, 1998; Djenoune et al., 2014; Jalalvand et al., 2014; Orts-Del’Immagine et al., 2014, 2017). dPKD2L1 cells have also a small round cell body (~10 μm diameter) and mostly present a bipolar morphology similar to that of CSF-cNs (Fig 2A1-5 and 3). Although neurites generally appeared less labelled against PKD2L1 than cell bodies, we often could detect dPKD2L1 cells with long PKD2L1^+^ neurites projecting in the ventro-dorsal axis along the midline and towards the CC (Fig. 2A). Therefore, to assess whether dPKD2L1^+^ cells may, alike CSF-cNs, contact the CC, we looked for dPKD2L1^+^ cell bodies with a neurite that could be followed all the way to the CC. In the 200 cervical and medullar sections that were analyzed, we could indeed clearly observe 10-15 of such cells with a long ventro-dorsal process (>70μm) that appears to contact the CC and to end with the typical bud (Fig. 2A1-5). These results indicate that dPKD2L1^+^ cells, at least some of them, might be in contact with the CSF. However, this number appears low, and one might wonder whether dPKD2L1^+^ cells represent a distal population of CSF-cNs. Nevertheless, in the CC region, we also observed numerous long ventral CC-contacting PKD2L1^+^ neurites (without cell bodies) that might belong to dPKD2L1 cells and were cut during the section preparation (not shown). We also observed on cervSC and Med sections a large number of dPKD2L1^+^ cells and neurites aligned along the midline axis and forming cell chains, so that isolated cell bodies and processes toward the CC could hardly be distinguished (Fig. 2B1–5 and 2C). Therefore, the number of distal CSF-cNs might be underestimated. Using the PKD2L1:EGFP mouse model, we confirm this observation and moreover indicate that dPKD2L1 cell chains (Fig. 2C, white arrows) were intermingled with GFP^+^ CSF-cNs axonal ventromedial projections (Fig. 2C, black arrows). Finally, whereas CSF-cNs exhibited only one dendritic process toward the CC, we also observed dPKD2L1 cells that exhibit a more complex organization with long processes projecting in various directions (multipolar morphology), even perpendicular to the ventro-medial axis, or in a direction opposite to the CC (Fig. 2B2, white arrows and see Fig. 3). These cells could also be observed as clusters (Fig. 2B3). These results would suggest that dPKD2L1 cells form two subpopulations: one largely bipolar and potentially in contact with the CC and one multipolar localized more distal and lateral from the CC and the midline, respectively. However, these observations would need further investigation to be confirmed.

**Fig. 3:**
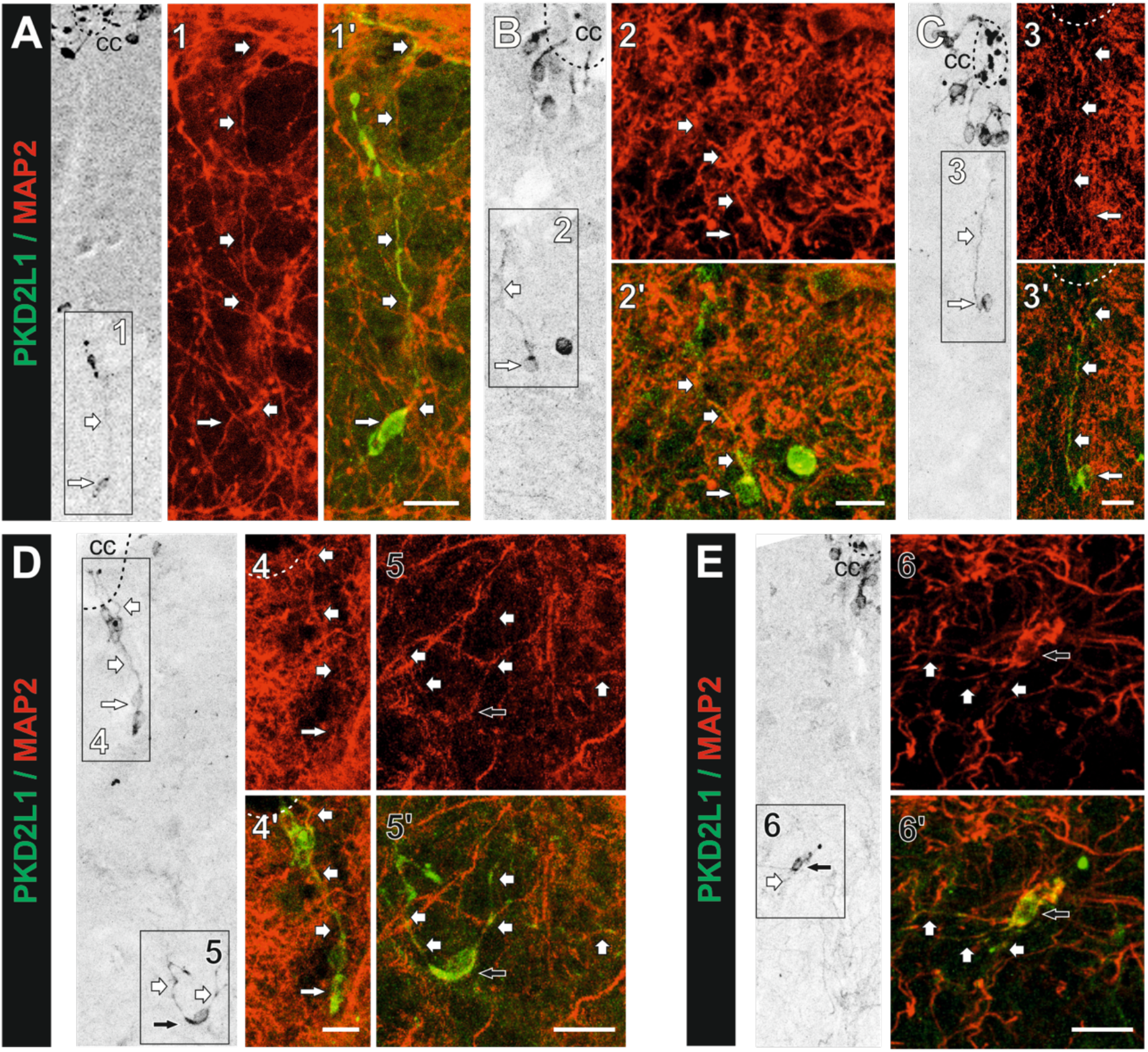
Identification of dPKD2L1 cell dendritic processes. MAP2/PKD2L1 immunolabelled cervSC sections showing examples of dPKD2L1 cell morphologies. For each examples the images in grey scale indicate the localization of the cells illustrated with images (numbered) at higher magnification with the MAP2 immunolabelling (red) and the PKD2L1/MAP2 merged staining (red/green). **A1, B2, C3** and **D4.** show bipolar cells extending a long dendrite (MAP2^+^) toward the CC (>100 μm from the CC) sometimes ending with a protrusion in the CC. **D5** and **E.** Examples of dPKD2L1 cells with multidirectional MAP2^+^ dendrites. Dashed lines: CC border. Thin white or black arrows: bi- and multipolar dPKD2L1 cells, respectively; Large white arrows: PKD2L1^+^/MAP2^+^ dendritic process. Scale bar: 20 μm; All images, Z-projections of 5 optical sections.

### Dendritic and axonal projections for dPKD2L1 cells

CSF-cNs have been shown to project a dendrite (MAP2 positive) to the CC, but the axon projection path remains an unresolved issue to date (Orts-Del’immagine et al., 2012; Orts-Del’Immagine et al., 2014). Here, in order to characterize the morphology of dPKD2L1 cells, we first performed MAP2/PKD2L1 dual immunolabelling to identify their dendritic projections. Our observations indicate that the long processes projecting from dPKD2L1 cells do co-express MAP2 and PKD2L1 and are therefore long dendrites extending to the CC (Fig. 3). Further, these MAP2^+^ processes reached the CC, where they end with a protrusion exhibiting a strong IR against PKD2L1 (Fig. 3A, B and D). We mentioned that some dPKD2L1 cells have multidirectional processes, perpendicular to the ventro-medial axis, or in a direction opposite to the CC (Fig. 1–3). Some of the ectopic multidirectional processes are immunolabeled against MAP2 as well as PKD2L1 and therefore represent dendritic projections (Fig. 3D5 and E). These multipolar dPKD2L1 cells were found in the most ventral zones, more lateral from the midline and their dendrites were not directed toward the CC. However, in PKD2L1^+^ processes a clear MAP2 immunolabelling was often hardly detectable because of the high density of MAP2^+^ processes in the parenchyma and since MAP2 IR was generally low in dPKD2L1 cells (probably due to their low mature phenotype, see below). Thus, we cannot exclude that some of these dPKD2L1^+^ processes may be axons rather than dendrites. Nevertheless, these results indicate that dPKD2L1 cells have a neuronal phenotype (see below) and will be denominated as neurons. Next, using immunohistofluorescence approaches in transgenic mouse models expressing either the tdTomato or GFP cytosolic markers (PKD-tdTomato and PKD-GFP mice), we addressed this question (Fig. 4). Our initial results, obtained in sagittal cervSC sections immunolabelled against PKD2L1, indicate the presence of long rostrocaudal PKD2L1^+^ fibers in the ventral region corresponding to the white matter (Fig. 4A-B, large white arrows). These fibers form longitudinal rostro-caudal projections presumably originating from CSF-cNs, as reported by Stoeckel and colleagues (2003), and potentially also from dPKD2L1 neurons. Note that fibers oriented in the dorso-ventral axis could be seen to join these fiber tracts (black arrows in Fig. 4B and see Fig. 4F) and that dPKD2L1^+^ cells bodies could be observed in close contact with these fibers (asterisk in Fig. 4A-C and E). To confirm this observation, we prepared SC tissue sections from PKD-tdTomato mice to visualize cell morphology and axonal projection from the tdTomato cytosolic fluorescence. In transverse cervSC sections, we confirmed, in agreement with Stoeckel and colleagues (2003), the presence of bilateral fiber bundles (patches of tdTomato fluorescence) in a zone corresponding to the ventral median fissure (Fig. 4C, white arrows). We also observed tdTomato^+^/PKD2L1^+^ cells as well as ventro-dorsal projections surrounding these tdTomato^+^ fiber bundles (Fig. 4C and D, asterisk and black arrows, respectively). However, we could not confirm that the axons express PKD2L1 since the tdTomato^+^ fiber bundles exhibited very low or no IR against PKD2L1 (Fig. 4C-E). Nevertheless, our data indicate that these tdTomato^+^ fibers correspond to axons since they are negative to immunolabelling against MAP2, a dendritic marker (Fig. 4 D). Our results were confirmed using PKD-GFP mice as illustrated in Figure 4E and indicate the presence of PKD2L1^+^/GFP^+^ neurites, presumably dendrites from dPKD2L1 neurons, running parallel to GFP^+^ fibers, presumably axons (Fig. 4E, white and black arrows). Further, using sagittal cervSC sections prepared from PKD-EGF mice, we could observe GFP^+^ dPKD2L1 neurons with a bipolar morphology and two neurites projecting in opposite direction along the dorsoventral axis. One runs dorsally towards the CC, presumably a dendrite, and the second projects ventrally, presumably axons, and joined the longitudinal rostro-caudal fibers bundle (Fig. 4F). Unfortunately, the axonal nature of the latter GFP positive projections could not be confirmed since, as we reported earlier (Orts-Del’Immagine et al., 2014), PKD2L1^+^ cells, have an intermediate maturity state (see also below), and do not express classical axonal markers such as neurofilament 160 kDa (NF160; Fig. 4 F) or neurofilament heavy polypeptide NF-H (200 kDa NF-H).

**Fig. 4:**
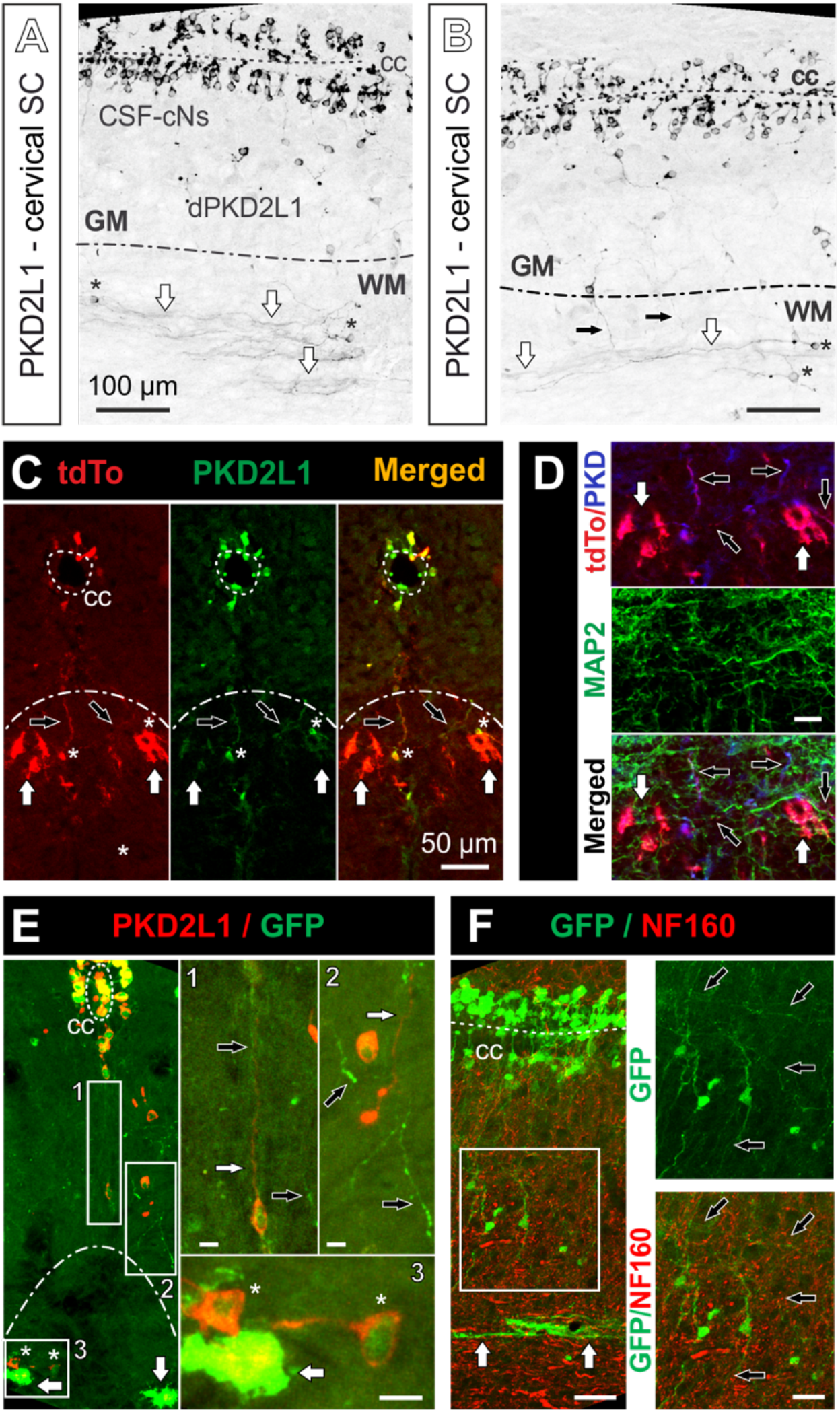
Axon from PKD2L1^+^ neurons regroup in longitudinal ventromedial fiber bundles. **A** and **B.** Examples of a representative PKD2L1 immunolabelled sagittal section (cervSC) showing dPKD2L1 cells localized between CC and grey matter (GM) and longitudinal neurites in the white matter (WM). Note the presence of dorso-ventral dPKD2L1^+^ neurite joining the longitudinal fiber bundles (black arrows) as well as dPKD2L1 cells with long rostrocaudal processes (asterisk). **C.** PKD2L1 immunolabelling in a PKD-tdTomato mice (cervSC coronal section) showing the presence PKD2L1^+^/tdTomato^+^ cells around the CC and in the vicinity (asterisk) of bilateral PKD2L1^−^/tdTomato^+^ ventral fiber bundles (white large arrows). Note the presence of dorso-ventral PKD2L1^+^/tdTomato^+^ neurites (black arrows). **D.** High Magnification of the region in C of the tdTomato^+^ ventral fiber bundles showing immunolabelling for PKD2DL1 (blue), MAP2 (green) and tdTomato fluorescence (red). Black arrows: PKD2L1^+^/tdTomato^+^/MAP2^+^ dendrites; Large white arrows: PKD2L1^−^/tdTomato^+^/MAP^−^ fiber bundles. **E.** PKD2L1/GFP immunolabelling in a PKD-GFP mouse (cervSC coronal section) showing bilateral PKD2L1^+^/GFP^+^ ventral axon bundles (large white arrow), PKD2L1^+^/GFP^+^ (thin white arrows, inset 1) and PKD2L1^−^/GFP^+^ (black arrows, inset 2) ventral processes. Inset 3 shows PKD2L1^+^/GFP^+^ cells (asterisks) contacting these ventral axon bundles. **F.** Example of NF160/GFP immunostaining in a sagittal section illustrating ventral rostro-caudal of PKD2L1^+^/GFP^+^ fibers bundles that are NF160^−^ (white arrows) as well as numerous bipolar GFP^+^dPKD2L1 cells with dorsal neurites directed toward the CC and ventral neurites (black arrows, inset). Scale bar: 100 μm (A, B, F); 50 μm (C), 20 μm (D) and 10 μm (E1-3); All images, Z-projections of 5 optical sections.

### Central canal organization and dPKD2L1 neurons distribution during development

PKD2L1^+^ CSF-cNs were shown to appear around the embryonic day 15 (E15) in mice and 18 somite stage in zebrafish larvae (Djenoune et al., 2014; Kutna et al., 2013; Petracca et al., 2016), however their origin, distribution and organization around and at distance from the CC is unclear. To address this question, we performed histological experiment in mice aged from E15 to 3M. We first analyzed the properties of the CC using PKD-tdTomato transgenic mice between E15 and E18 as well as P0 and 3M (Fig. 5A-C). In E15 embryo cervSC transverse sections, the CC appears as a long slit of ~250 μm length (Fig. 5B) and surrounded by the neuroepithelial layer (Shibata et al., 1997; Kutna et al., 2013). With development, the CC length progressively decreases to reach its final length at P0 (Fig. 5A and B; E15: 268 ± 32 μm; E16: 72 ± 11 μm; E18: 46 ± 9 μm; P0: 36 ± 8 μm and 3M: 36 ± 8 μm; One-Way ANOVA: F =2141.3, dF = 4, p(F) < 2.2.10^−16^). Over time, the CC decreases in length presumably with the fusion of its ventral part (Sturrock, 1981) and is localized in a more dorsal position of the SC. The CC progressively acquires a circular shape, and its ventral edge is more distant from the SC ventral pole (Fig. 8A and C; E15: 155 ± 22 μm; E16: 206 ± 23 μm; E18: 271 ± 38 μm; P0: 446 ± 87 μm and 3M: 822 ± 92 μm; One-Way ANOVA: F =1430.5, dF = 4, p(F) < 2.2.10^−16^).

**Fig. 5:**
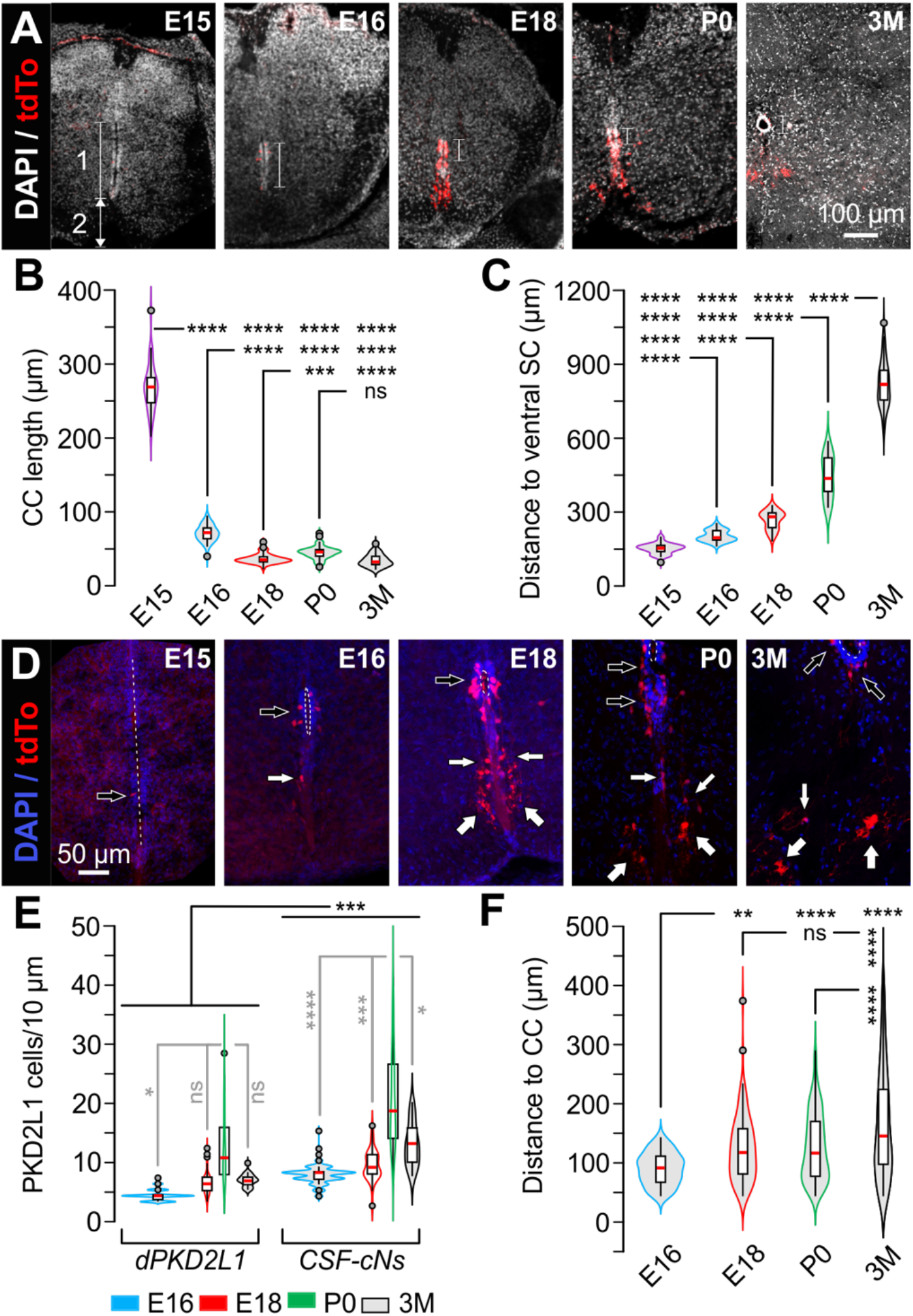
Central canal morphology and dPKD2L1 cells organization during development. **A.** cervSC sections (DAPI staining) from a PKD-tdTomato mouse illustrating central canal evolution and the distribution of dTomaoto^+^ (red) cells at different 3 developmental stages (E15, E16, E18), postnatal (P0) and adult ages (3M). The white bar (1) and the white double arrow (2) represents the CC length and the distance between CC edge and ventral SC, respectively. **B** and **C.** Summary whiskers box plots with violin plots (data point density) comparing the CC length (B) and the distance between the CC ventral edge and the SC ventral pole (C). during the 4 analyzed time points. **D.** Magnification images of the CC region at the 4 analyzed time points showing the distribution of tdTomato^+^ cells around the CC and in the ventral region. Black arrows: tdTomato^+^ cell bodies around the CC; White arrows: distal tdTomato^+^ cells; Large white arrows: tdTomato^+^ axons. **E.** Summary whiskers box plots with violin plots (data point density) of dPKD2L1 neurons and CSF-cN number between E16 and 3M. **F.** Summary whiskers box plots with violin plots (data point density) of distal tdTomato^+^ cell distance from the CC in the ventral SC between E16 and 3M. Scale bar: 100 μm (A) and 50 μm (D); Z-projections of 10 (A) and 6 (D, E15), 10 (D, E16 and E18) 12 (D, P0), 15 (D, 3M) optical sections.

Next, we analyzed at the same ages, the number and distribution of tdTomato^+^ cells (*i.e.* PKD2L1 cells) in cervSC sections prepared from PKD-tdTomato mice. When looking at E15, we only found rare tdTomato^+^ cells (less than one per section) that are generally localized in the ventral half of the CC (Fig. 5D, black arrow). Note that because of the low number of tdTomato^+^ cells at E15, it was not possible to have a quantitative analysis. At E16, we counted 6 ± 2 tdTomato^+^ cells mainly distributed around the CC (Fig. 5D and E; 5 ± 2 and 1 ± 1 CSF-cNs and dPKDL21 cells, respectively). With development the number of tdTomato^+^ cells progressively increases to reach a peak at around P0. On average, the number of CSF-cNs was higher than that of dPKD2L1 neurons at all ages and we found: 4.8 ± 1.9 (E16), 6.8 ± 2.5 (E18), 17.9 ± 7.9 (P0) and 10.3 ± 3.6 (3M) CSF-cNs/10 μm of tissue *vs.* 1.1 ± 0.9 (E16), 3.2 ± 1.8 (E18), 8.6 ± 5.5 (P0) and 3.6 ± 1.0 (3M) dPKD2L1 cells/10 μm of tissue (Fig. 5D and E; *nlme* model: χ^2^(Type) = 375.71, dF = 1, p(χ^2^) < 2.2.10^−16^). Although, dPKD2L1 neurons are less numerous than CSF-cNs at all ages (Fig. 8E; *nlme* model: χ^2^(Age) = 50.186, dF = 3, p(χ^2^) = 7.292.10^−11^), the average percentage of dPKD2L1 neurons compared to the total PKD2L1^+^ cell population increases with development to reach a peak between E18 and P0 (E16: 18 ± 13%; E18: 32 ± 16%; P0: 30 ± 10% and 3M: 27 ± 8%; One-Way ANOVA: F(Age) = 22.488, dF = 3, p(F)= 4.092.10^−13^). Further, we calculated the ratio of dPKD2L1 neurons compared to CSF-cNs and found that it first increases from E16 up to P0 before decreasing at 3M (dPKD2L1/CSF-cN: 25 ± 24% at E16; 47 ± 23% at E18; 46 ± 20% at P0 and 38 ± 16% at 3M; One-Way ANOVA: F = 21.094, dF = 3; p(F) = 1.292.10-^12^). These results indicate that new PKD2L1 expressing neurons might be generated up to early postnatal days and that dPKD2L1 neurons may migrate from the CC region to more ventral ones. Finally, this increase in dPKD2L1 neuron number during development was accompanied by a progressive increase in their distance to the CC: 92 ± 27 μm at E16, 125 ± 56 μm at E18, 129 ± 57 μm at P0 and 168 ± 92 μm at 3M (Fig. 8F; One-Way ANOVA: F = 28.611, dF = 3, p(F) < 2.2.10^−16^). In parallel, we observed that a proportion of dPKD2L1 appear to migrate laterally from the midline since the average distance increased with age, from 15.2 ± 10.7 μm at E16 to 42.9 ± 53.6 at 3M (One-Way ANOVA: F =21.513, dF = 4, p(F) = 3.976.10^−13^). Our data further suggest that the axons of CSF-cNs and probably dPKD2L1 neurons progressively grow ventrally to join the region of the median fissure where they regroup in bilateral fiber bundles (Fig. 5D, large white arrows and see Fig. 4).

### Postnatal dPKD2L1 neurons are immature and are present within the ependymal niche

To further specify the phenotype and cell environment of these dPKD2L1 neurons, we analyzed their organization and properties in neonatal mice (P0-2). Our data indicate that the number of dPKD2L1 neurons and CSF-cNs reaches a peak at P0 and was higher compared to tissue from 3M both at the cervSC and *Medulla* levels (CSF-cNs: 17.9 ± 7.9 *vs.* 10.3 ± 3.6 in the cervSC and 19.4 ± 7.1 *vs.* 9.8 ± 3.5 in the *Medulla* at P0 and 3M, respectively and dPKD2L1 neurons: 8.6 ± 5.5 *vs.* 3.6 ± 1.5 in the cervSC and 8.2 ± 3.4 *vs.* 3.3 ± 1.5 in the *Medulla* at P0 and 3M, respectively and see above Fig. 5E; *nlme* model χ^2^(Age) = 1.5611, dF = 1, p(χ^2^) = 0.2189 and χ^2^ (Region) = 294.5257, dF = 3, p(χ^2^) < 2.2.10^−16^ with an interaction Age:Region: χ^2^ = 11.8594, dF = 3, p(χ^2^) = 0.0078). Further, at P0, dPKD2L1 neurons appear closer to the CC than in tissue from 3M mice and restrained to the grey matter, as the white matter is almost absent in the SC at this stage (Fig. 6A, insert). However, numerous dPKD2L1 neurons very distant from the CC (>250 μm) could also be observed at P0, especially in the more rostral region (see Fig. 5F). At both ages, the distance from the CC of dPKD2L1 neuron increases in the caudo-rostral axis and dPKD2L1 are the most distant in the spMed compare to cervSC (129 ± 57 μm, 129 ± 46 and 171 ± 101 at the cervSC, c and spMED at P0 and 168 ± 92 μm, 166 ± 85 and 239 ± 109 at the cervSC, c and spMED at 3M; *nlme* model χ^2^(Age) = 46.086, dF = 1, p(χ^2^) = 1.132.10^−11^ and χ^2^ (Region) = 77.113, dF = 2, p(χ^2^) < 2.2.10^−16^ with an interaction Age:Region: χ^2^ = 11.107, dF = 2, p(χ^2^) = 0.0038). In a previous report, we indicated that CSF-cNs exhibited an immature phenotype at early postnatal stages. We therefore performed dual immunolabelling against PKD2L1 and NeuN or HuC/D to assess whether distal neurons would share this feature at P0. Our results show that in the cervSC and Med, the ventro-medial area is devoid of NeuN^+^ (Fig. 6A) or HuC/D^+^ neurons (Fig. 6C) and that dPKD2L1 neurons do not express NeuN. Next, we looked for the organization and projection of ependymal cells at P0 using immunolabelling against vimentin (Vim), a selective marker. We observed Vim^+^ cells around the CC as well as long Vim^+^ fibers projecting ventrally and forming two parallel bundles within an area devoid of NeuN and HuC/D immunostaining that would correspond to the niche formed by radial ependyma cells (Fig. 6A-C). Interestingly, most dPKD2L1 neurons are aligned at the border of the neuronal zone (NeuN^+^, cervSC in Fig. 6A or HuC/D^+^, spMed in Fig. 6A) along or within the 2 ependymal fiber tracts (Fig. 6B). In this zone dPKD2L1 neurons often appear forming ventro-medial cell chains (Fig. 6C). Moreover, at P0, we observed a dorso-ventral zone along the midline that exhibits strong PSA-NCAM IR where numerous dPKD2L1 neurons are localized (Fig. 6D). Moreover, at higher magnification dPKD2L1 neurons, even distant from the midline, are immunoreactive against PSA-NCAM (Fig. 6E and see Fig. 7). Finally, alike CSF-cNs, dPKD2L1 neurons express DCX (Fig. 6F and see Fig. 7).

**Fig. 6:**
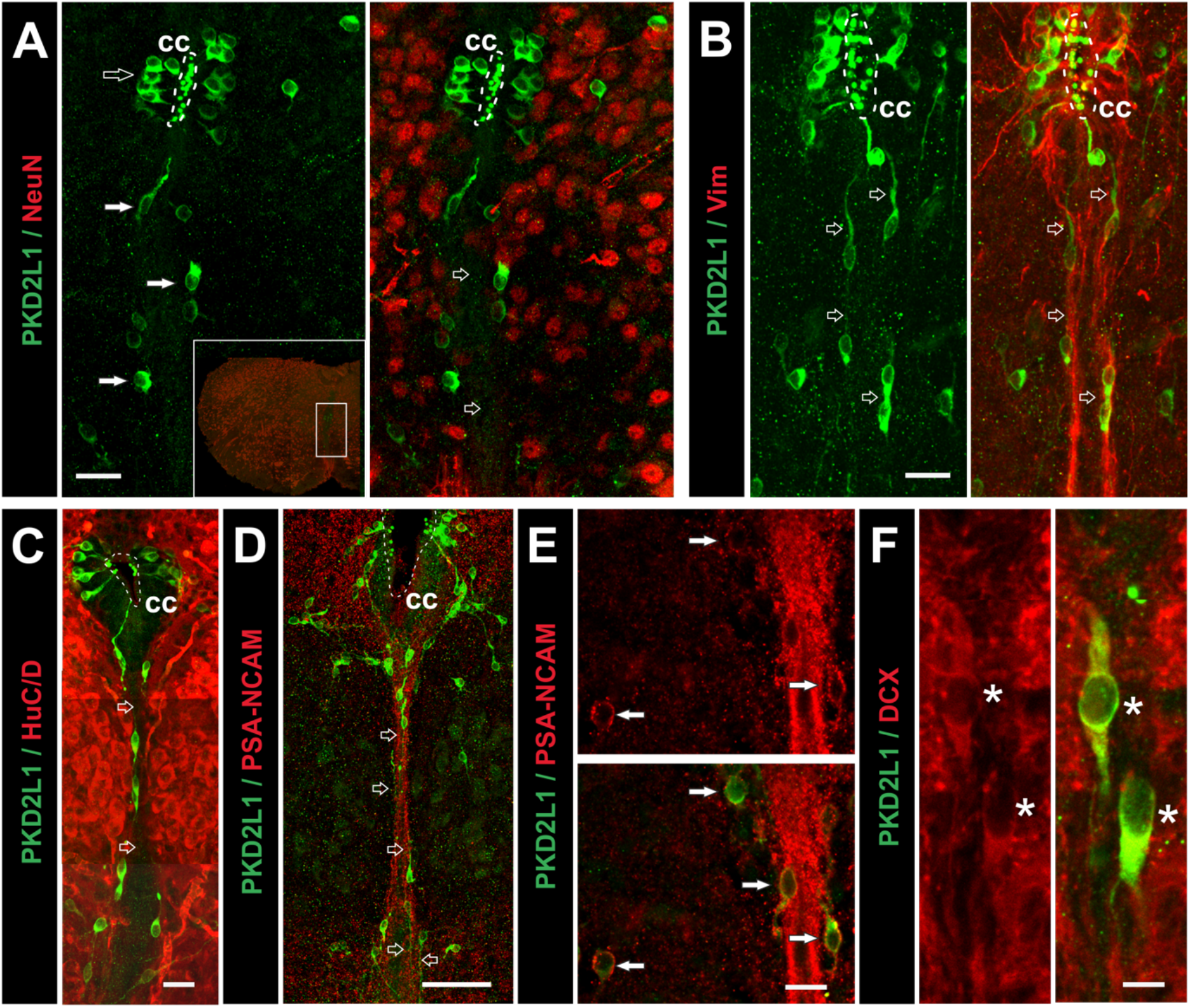
Postnatal dPKD2L1 neurons are immature and distributed in the ependymal niche. **A.** PKD2L1/NeuN immunolabelling in a cervSC section prepared from a P0 mouse and showing the absence of NeuN^+^ (merged, red) cells in a ventro-medial lane and the presence of 2 rows of dPKD2L1 cells (green, white arrows) and processes (black large arrows) at each border. Inset: cervSC section showing the illustrated area and the zone magnified. **B.** PKD2L1/Vim immunolabelling in a cervSC section at P0 showing dPKD2L1 neurons and processes (green, black arrows) along the 2 dorso-ventral Vim^+^ fiber tracts (red). **C.** PKD2L1/HuC/D immunolabelling (Med section) at P0 showing the ventro-medial midline zone devoid of HuC/D^+^ cells but the presence of dPKD2L1 cells and processes at its border. **D.** PSA-NCAM/PKD2L1 dual immunolabelling (P0 spMed section) showing dPKD2L1 cells and processes (black arrows) along the 2 main PSA-NCAM^+^ fiber tracts. **E.** Magnification of the central midline zone and PSA-NCAM/PKD2L1 dual immunolabelling (P0 spMed section) showing PSA-NCAM IR also in ventro-medial fibers but also in dPKD2L1 cells (white arrows). Note the close proximity between some dPKD2L1 cells and PSA-NCAM^+^ ependymal fibers. **F.** PKD2L1/DCX dual immunolabelling (P0 mice cervSC, ventro-medial region) showing the presence of DCX in dPKD2L1 cells (asterisk) but also in the surrounding grey matter. Scale bars: 20 μm (A, B, C, F); 50 μm (D); 20 μm (E); Z-projections of 13,15 (A-D) and 3-5 (E-F) optical sections.

**Fig. 7:**
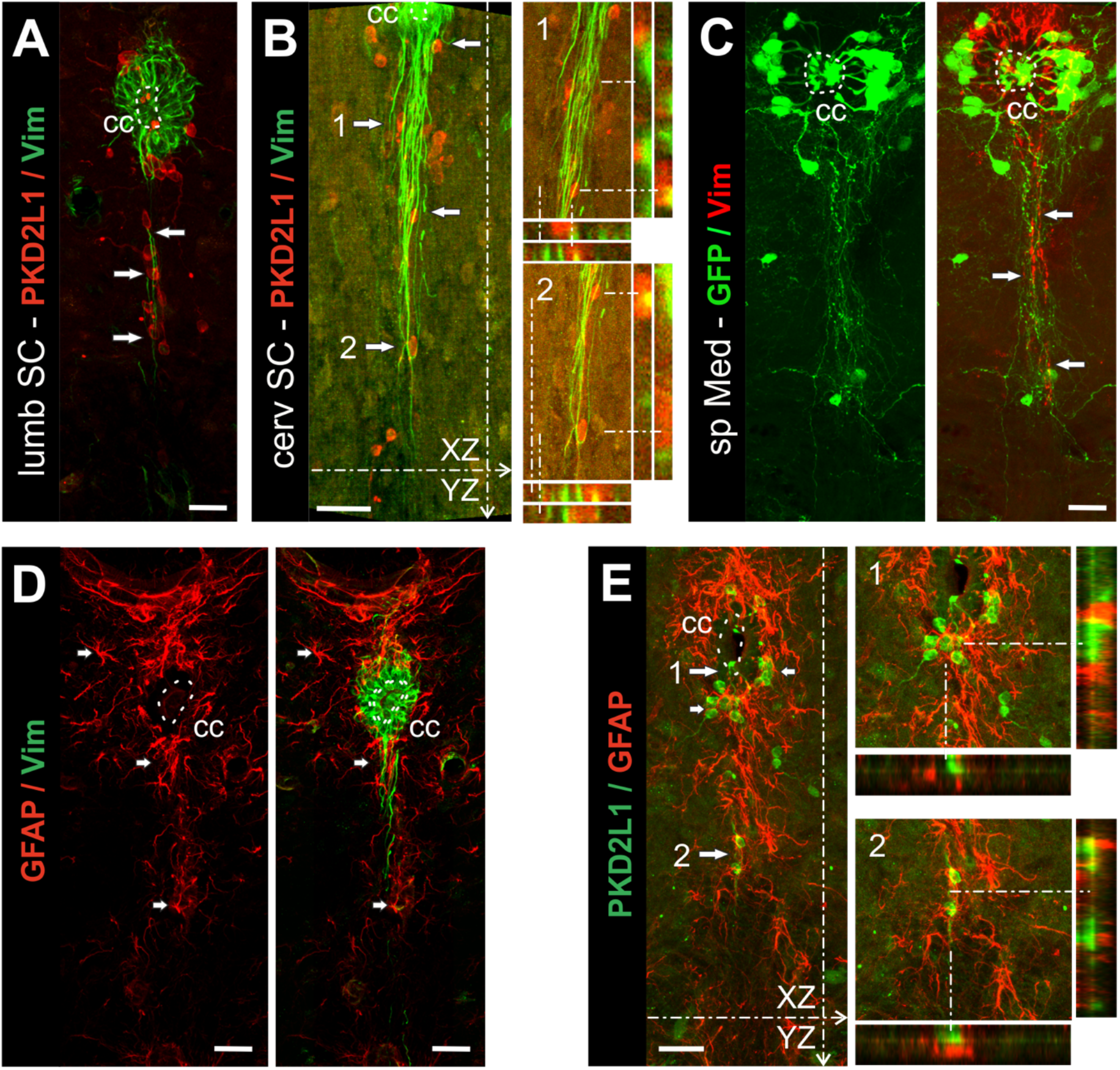
In adult mice, dPKD2L1 cells are constitutive of a ventro-medial ependymal cell niche. **A** and **B.** Vimentin (Vim)/PKD2L1 immunolabelling in lumbSC and cervSC sections, respectively showing dPKD2L1 cells and processes aligned along the ependymal fibers. **B.** Inset 1 and 2 represent orthogonal projections from B at 2 levels (dashed lines) illustrating the close apposition of Vim^+^ fibers around dPKD2L1 somas. **C.** GFP/Vim dual labelling in a spMed section (PKD-GFP mouse) showing that PKD2L1/GFP^+^ axons and dPKD2L1 cells (white arrows) are intermingled with ependymal fibers (red). **D.** Vim/GFAP dual immunolabelling in a cervSC section showing a high density of GFAP^+^ astrocytes (red, white arrows) in the subependymal layer and along the ventro-medial Vim^+^ ependymal fibers (green). **E.** PKD2L1/GFAP dual immunolabelling in a cervSC section showing colocalization of GFAP^+^ astrocytes (red) and dPKD2L1 cells (green, arrows) around the ependymal layer and in the ventro-medial region. Inset 1 and 2 represent orthogonal projections of E at 2 levels (dashed lines) showing the close apposition of GFAP processes around CSF-cNs (Top) and dPKD2L1 cells (Bottom). Scale bar: 25 μm; Z-projections of 10 (A-E), 5 (B1, 2; E1, 2) optical sections.

### dPKD2L1 cells and processes are constitutive of an ependymal/astroglial niche

We reported that with aging, CSF-cNs largely retained immature features, are in close contact with ependymal cells so that they were suggested to be part of the medullo-spinal stem cell niche. Here, we indicate that at P0 dPKD2L1 neurons are mainly localized in the ventro-medial region close to Vim^+^ and PSA-NCAM^+^ fibers. However, to date, no data are available regarding their environment and their interaction within the stem cells niche at later stage. To elucidate this point, we conducted immunohistofluorescence experiments to analyze dPKD2L1 cell organization within the EZ. On transverse lumbSC and cervSC sections at 3M, immunolabelling against Vim, indicates that ependymal cells around the CC also project long Vim^+^ fibers to the ventral region and that these fibers form dorso-ventral fiber bundles along the midline (Fig. 7A-B). When looking, in the adult lumbSC and cervSC sections, at the localization of dPKD2L1 neurons in respect to Vim^+^ fibers, we observed that the dPKD2L1 neurons present along the midline are found in apposition or close to Vim^+^ radial ependymal fibers. Further, their processes generally follow the trajectory of the Vim^+^ ependymal fibers (Fig. 7A-C). Our analysis using orthogonal projections strongly suggest potential contacts between Vim^+^ fibers and dPKD2L1 cell bodies (Fig. 7B1 and 2). Note that dPKD2L1 localized in more lateral zone from the midline do not contact the ependymal fibers. As mentioned above, the PKD2L1:GFP mice model allows visualizing PKD2L1 cell morphology but also axonal projection. When comparing the distribution of GFP^+^ projections with that of Vim^+^ fibers, we noticed that GFP^+^ ventro-medial axons and cell bodies occupy the same territories and follow the same trajectory than ependymal fibers (Fig. 7C). One might therefore suggest that CSF-cNs and dPK2L1 cells within the ependymal niche would interact with ependymal cells and fibers. Previous reports indicated that the EZ and the region along the ventro-medial region is enriched with astrocytes (Hamilton et al., 2009; Hugnot and Franzen, 2011). Here, our results, using double immunolabelling against Vimentin and GFAP (a marker for astrocytes), reveal a high density of GFAP staining around the EZ as well along of the ventro-medial Vim^+^ fibers (Fig. 7D). To assess possible interaction between PKD2L1^+^ neurons and astrocytes, we performed dual immunostaining experiments using antibodies against PKD2L1 and GFAP. Our data indicates that CSF-cNs but also dPKD2L1 neurons are present within an astrocyte enriched zone, are surrounded with GFAP+ processes that appear to cover CSF-cNs and dPKD2L1 neuron somas (orthogonal projection in Fig. 7E).

### dPKD2L1 cells have the same phenotypical determinants as CSF-cNs in adult

We indicated that CSF-cNs and dPKD2L1 neurons present similar immature phenotype at birth. Further, in adult mice, CSF-cNs exhibit the unique feature to retain an incomplete maturity state, as indicated by low NeuN immunostaining and by the presence of some immaturity markers as doublecortine (DCX) and the pattern determinant Nkx6.1 (Sabourin et al., 2009; Orts-Del’Immagine et al., 2014, 2017). Here, in order to compare dPKD2L1 neurons with CSF-cNs in adult mice (3M) and assess whether they also retain similar properties, we analyzed the expression of various neuronal cell markers. First, we conducted immunolabelling against GABA and our results indicate that dPKD2L1 cells, alike CSF-cNs (Djenoune et al., 2014; Orts-Del’Immagine et al., 2014), are GABAergic. Note that we observed GABA^+^ cells located in the ventro-medial white matter (Fig. 8A). Using GAD67/PKD2L1 double immunolabelling, we show that the isoform 67 of the glutamic acid decarboxylase (GAD67), the enzyme responsible for GABA production, is also detected in dPKD2L1 cells (Fig. 8B, white arrows), although clear detection was generally difficult due to the widespread expression of GAD in the parenchyma (Fig. 8B). Next, PKD2L1/NeuN immunolabelling shows that NeuN IR intensity is very low in dPKD2L1 cells (by comparison to the surrounding neurons), some of them being clearly devoid of NeuN labelling (Fig. 8C, white arrows). Nevertheless, on average 63 ± 8% and 69 ± 13% of dPKD2L1 neurons at the cervSC and cMed levels express NeuN while this percentage reached 89 ± 9% at the spMed level (2-3 animals; *nlme* Model: χ^2^(Region) = 35.732, dF = 2, p(χ^2^) = 1.74.10^−8^). When analyzing the percentage of NeuN^+^ CSF-cNs, we found a similar distribution (cervSc: 54 ± 12%, cMed: 77 ± 2% and spMed: 92 ± 7%; χ^2^ (Type) = 0.015, dF = 1, p = 0.9014). Moreover, whereas NeuN^+^ neurons were all present within the grey matter (so that NeuN immunostaining permit to clearly distinguish the grey matter from the white matter), most dPKD2L1 neurons were usually present at the border of the grey/white matter or within the white matter (Fig. 8C). Similar observations were obtained using the neuronal marker Hu/CD (not shown). On the other hand, analysis of DCX/PKD2L1 immunolabelling in adult mice shows that dPKD2L1 cells are generally DCX^+^ (~90%) (Fig. 8D) with an expression level constant along the caudo-rostral axis (cervSC: 90 ± 9%, cMed: 85 ± 8% and spMed: 89 ± 9%; 2-3 animals; *nlme* Model: χ^2^(Region) = 0.844, dF = 2, p = 0.6557). Next, CSF-cNs were reported to express Nkx6.1, a homeobox transcription factor involved in ventral neural patterning, even in 1-year-old mice. The analysis of Nkx6.1 immunolabelling indicate that dPKD2L1 cells do express nuclear Nkx6.1 and that dPKD2L1 cells were the sole Nkx6.1^+^ cells detected in the whole adult SC parenchyma (Fig. 8E). Finally, concerning PSA-NCAM, another marker for juvenile neurons, we previously reported that PSA-NCAM IR was not detected in CSF-cNs in adult mice but rather in the ependymal cells (Chatelin et al., 2001; Sabourin et al., 2009; Orts-Del’Immagine et al., 2017). Here, analysis of PSA-NCAM/PKD2L1 dual immunolabelling in the ventral SC of adult mice confirms the results obtained in P0 mice with the presence of PSA-CAM around the CC in ependymal cell bodies as well as in long medial fibers emerging from the CC region. However, in contrast to the situation observed at P0, dPKD2L1 do not express anymore PSA-NCAM (Fig. 8F). Interestingly, we often found dPKD2L1^+^ cell bodies in close apposition with these PSA-NCAM^+^ long dorso-ventral fibers (Fig. 8F, arrows). Our dual Vimentin (Vim)/PSA-NCAM immunolabelling confirms that the PSA-NCAM^+^ fibers correspond to projections of ependymal cells from the CC along the midline as they were also Vim^+^ (Fig. 8G).

**Fig. 8:**
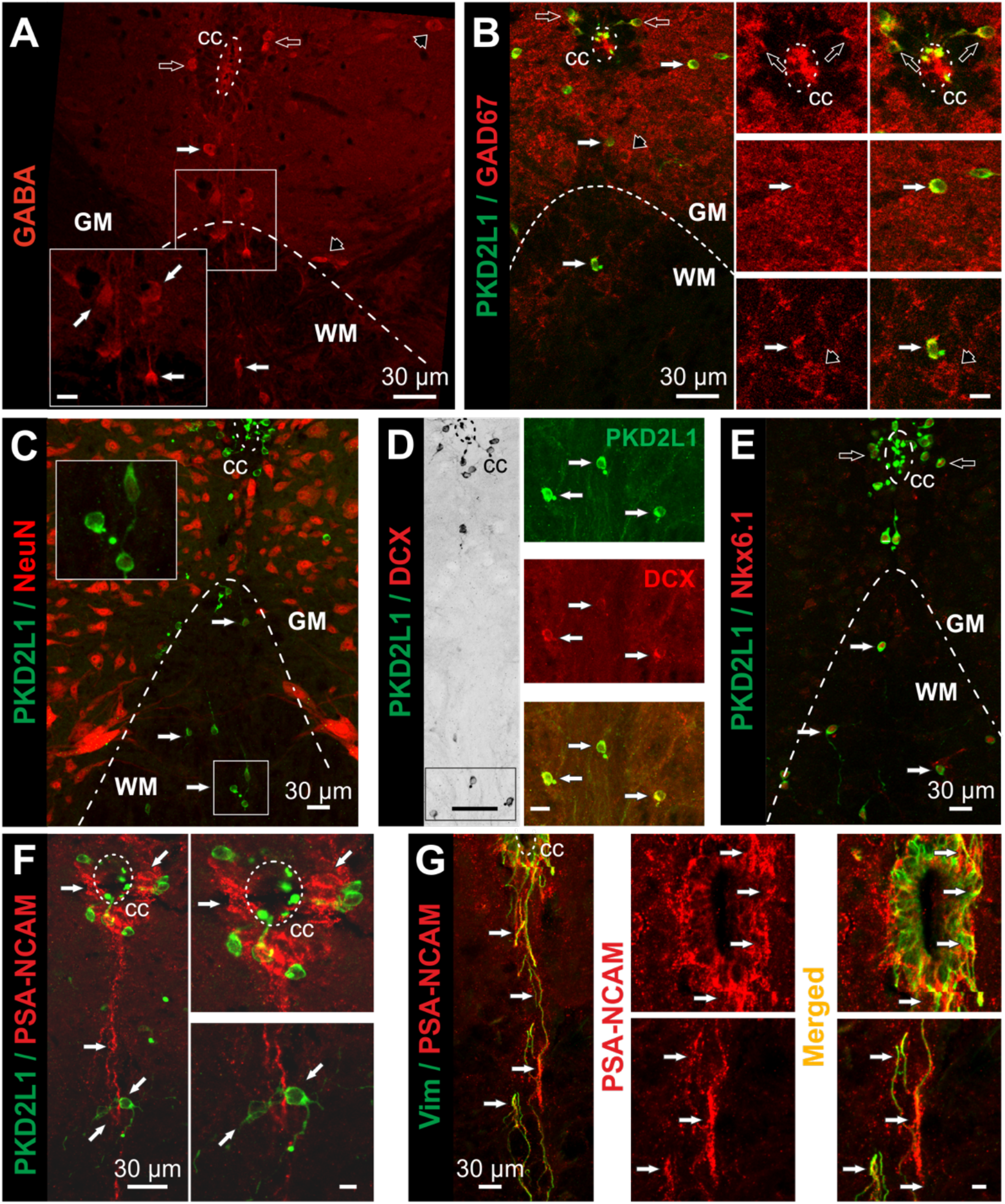
dPKD2L1 cells are GABAergic and present an immature phenotype in adult mice. **A.** Representative image showing GABA immunolabelling with GABA^+^ neurons (open arrows) around the CC and in the ventro-medial region (white arrows) that can be visualized. **B.** GAD67/PKD2L1 dual immunolabelling showing GAD67 IR in CSF-cNs (open arrows) and dPKD2L1 neurons (white arrows). Insets: magnification of the cells visualized in B showing GAD67^+^ (Left) and GAD67^+^/PKD2L1^+^ (Right, merged) cells around the CC (open arrows) and dPKD2L1 neurons. Black arrow points to a GAD67^+^ but PKD2L1^−^ neuron in B and in inset. **C.** NeuN/ PKD2L1 dual immunolabelling showing NeuN labelling in dPKD2L1 and CSF-cNs cells. Note that the NeuN IR is low compared to other neurons in the parenchyma and the presence of dPKD2L1 cells in the white matter (WM) devoid of NeuN IR. (higher magnification of 3 cells in the inset). **D.** Image (grey scale, Left) showing the localization of 3 dPKD2L1 neurons illustrated at higher magnification on the Right. Inset illustrate PKD2L1 (Top, arrows) and DCX (Middle) immunolabelling as well as the superposition (Bottom). **E.** Nkx6.1/PKD2L1 dual immunolabelling showing that CSF-cNs and dPKD2L1 neurons exhibit strong nuclear Nkx6.1 IR. **F.** PSA-NCAM/PKD2L1 dual showing PSA-NCAM IR around the CC and along ventro-medial fibers and that dPKD2L1 neurons and CSF-cNs are PSA-NCAM^−^. Inset magnification of the image on the Left in the CC (Top) and the ventral (Bottom) regions. Note the close proximity between dPKD2L1 neuron and PSA-NCAM^+^ ventral fibers. **G.** PSA-NCAM/Vimentin (Vim) dual immunolabelling showing a PSA-NCAM/Vim colocalisation around the CC and in the ventro-medial ependymal fibers. Coronal sections from the cervSC; Dotted-Dashed lines: limit between the grey (GM) and white matter (WM); Dashed lines: CC border; in A-E: Open arrows, CSF-cNs and White arrows, dPKD2L1^+^ cells; in F and G: ependymal cells and fibers. Black arrows in A and B: PKD2L1^−^ neurons in the parenchyma. Scale bar: 30 μm except D, Left: 50 μm; Z-projections of 2 (A), 3 (B), 10 (C), 3 (D), 10 (E), 10 (F and G) and 4-6 (insets in F and G) optical sections.

### dPKD2L1 cells exhibit functional properties similar to CSF-cNs

CSF-cNs exhibit characteristic functional properties and in particular present spontaneous PKD2L1 unitary current activity (Orts-Del’immagine et al., 2012; Jalalvand et al., 2016a, 2016b; Orts-Del’Immagine et al., 2016). To date the functional properties of dPKD2L1 neurons have never been investigated. In this last set of experiments, we therefore analyzed the electrophysiological properties of dPKD2L1^+^ cells in brainstem slices. Using acute slices prepared form PKD2L1:EGFP mice, we preformed whole-cell patch clamp recordings from GFP^+^ neurons located in ventro-medial *Medulla*. Figures 9 illustrates recordings from 2 GFP^+^ neurons present at 175 (Fig. 9A-B) and 250 μm (Fig. 9C-D) in the ventral region. All neurons recorded in the ventral region exhibit a low membrane capacitance (6.8 ± 3.7 pF, n = 7) and a high input resistance (2.3 ± 1.2 GΩ, n = 7) with a resting membrane potential (RMP) at −44.9 ± 1.2 mV (n = 7). When recorded in current-clamp mode, spontaneous action potential (AP) discharge activity can be recorded from the RMP (Fig. 9B_1_ and D_1_). To elicit APs, we injected DC current with à 500 ms pulse at +30 pA and either train of APs (Fig. 9B_1_, inset; n = 5) or a single AP (Fig. 9D_1_, inset; n = 2) were elicited. Typically, the AP threshold was at - 22.9 ± 0.7 mV and AP had an amplitude and a half width of 64.1 ± 5.2 mV and 3.9 ± 0.8 ms, respectively. In dPKD2L1^+^ neurons, where tonic firing was elicited, the average frequency of AP discharge was 20.2 ± 5.1 Hz (n = 5). Finally, as previously reported for CSF-cNs, PKD2L1 channels represent a hallmark for this unique neuronal population (Orts-Del’immagine et al., 2012; Orts-Del’Immagine et al., 2016), we therefore probed whether PKD2L1 channel activity could also be recorded in dPKD2L1 neurons. Our recordings performed in voltage-clamp mode at a holding potential set at −70 mV show that inward unitary current with the features of PKD2L1 channel are indeed observed in all dPKD2L1 neurons tested (Fig. 9B_2_ and D_2_). The average current amplitude was −13.0 ± 1.1 pA (n = 7) and the average open probability (N.Po) was 0.006 ± 0.004 (n = 7). In addition to single-channel current, inward currents with fast onset and exponential recovery, corresponding to synaptic currents, were observed in the recorded dPKD2L1^+^ neurons in the absence of antagonist against ionotropic GABAergic and glutamatergic receptors (Fig. 9B_2_ and D_2_). Note that we did not observe difference in their functional properties as a function of their distance to the CC.

**Fig. 9:**
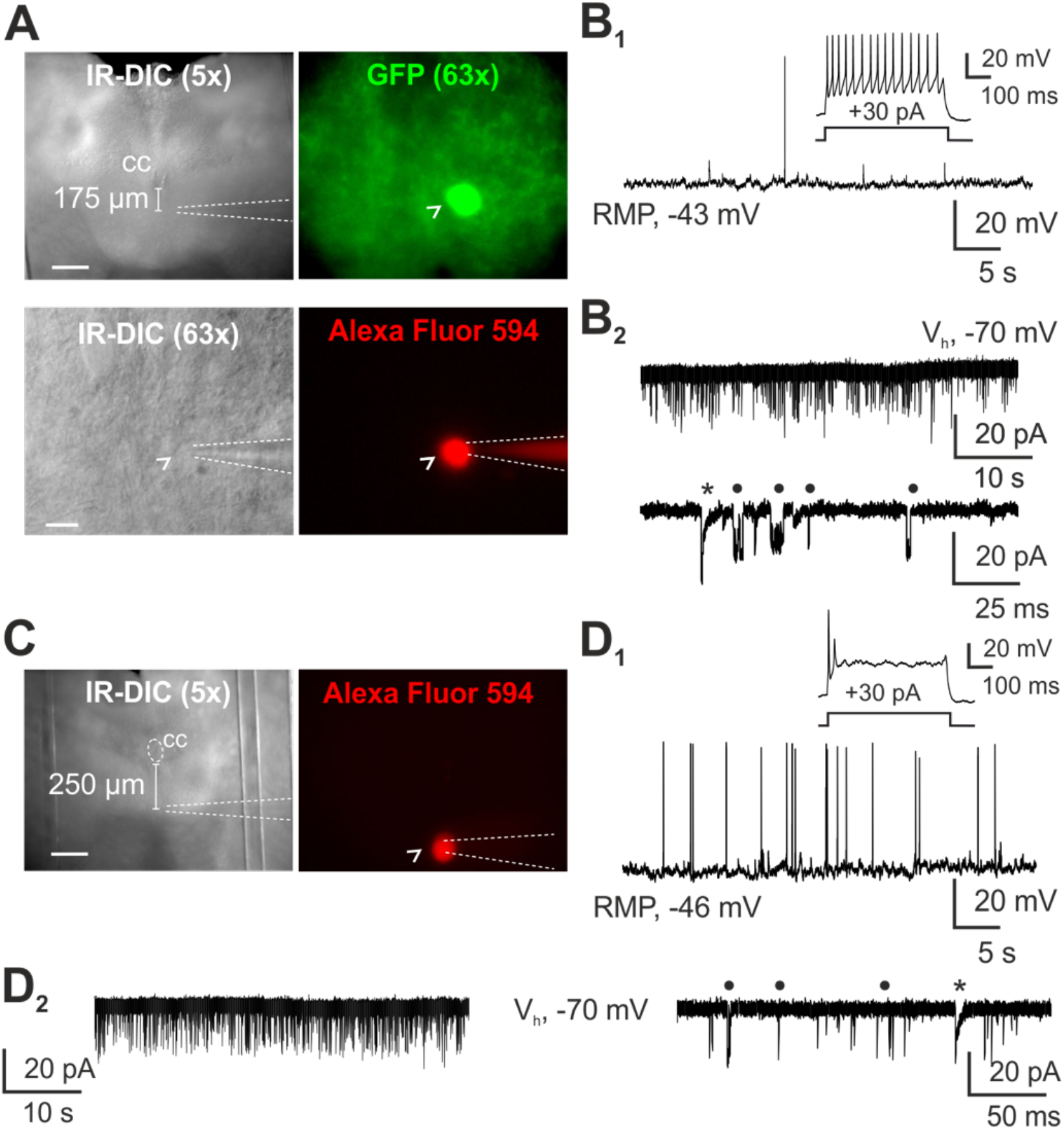
dPKD2L1 neurons exhibit spontaneous synaptic and unitary PKD2L1 current activities. **A** and **C.** Micrographs showing representative dPKD2L1 neuron expressing GFP (open arrowhead) located at 175 μm and 250 μm from the CC, respectively (from PKD-GFP mice). Recorded neurons are dialyzed with AlexaFluor 594 (red). **B** and **D.** Representative recordings from the cells shown in A and C, respectively. In current-clamp mode (**B_1_** and **D_1_**), spontaneous AP discharge activity recorded at the RMP. The insets illustrate the AP discharge pattern upon positive current injection. In voltage-clamp mode (**B_2_** and **D_2_**) and a holding potential of −70 mV, cells exhibit spontaneous inward currents (top trace). At slower time resolution, the same recordings show fast synaptic current (star) and characteristic PKD2L1 unitary current activity (circle). White dashed lines: position of the patch pipette. Scale bar is 200 μm for 5x magnification, and 10 μm for 63x magnification.

## DISCUSSION

Here, we describe the presence of distal PKD2L1 expressing cells (dPKD2L1 cells) localized along the medullo-spinal axis in the ventral region away from the CC. This zone is enriched with astrocytes and is formed by long dorso-ventral fibers originating from ependymal cells. dPKD2L1 cells are neurons and exhibit both phenotypically and functionally properties similar to those previously described for PKD2L1^+^ CSF-cNs around the CC. They are GABAergic and have an intermediate neuronal maturation state (low or no NeuN expression but DCX IR) that is retained in adult mice. Along the midline beneath the CC in the dorso-ventral ependymal/astroglial niche dPKD2L1 cells have a bipolar morphology with a long dendrite projecting toward the CC. We also observe dPKD2L1 cells with a more complex morphology that do not appear to send a projection to the CC and are localized in more lateral position from the midline. Finally, our results indicate that dPKD2L1 neurons appear at late embryonic stages and acquire during development a position distant from the CC and the midline while they lose the expression of PSA-NCAM. This suggest dPKD2L1 neurons may possess some migratory capabilities and move ventrally and laterally from the CC during early postnatal stages.

### A new population of PKD2L1 expressing neurons in the ventral spinal cord and Medulla

In our previous studies (Orts-Del’Immagine et al., 2014, 2017), we described the properties of PKD2L1^+^ neurons around the CC, designated here as CSF-cNs (Marichal et al., 2009; Djenoune et al., 2014, 2017; Orts-Del’Immagine et al., 2014, 2017; Petracca et al., 2016). Here we indicate that neurons expressing PKD2L1, dPKD2L1 cells, can be found in more ventral region (> 50 μm and up to several hundreds of microns) and are distributed along the medullo-spinal axis (from the lumbSC to the spMed). These cells, with small round cell bodies, are localized at distance from the CC in the ventral region as far as within the white matter. Interestingly, they are the only neurons present in this region. dPKD2L1 neurons are often observed as chain or cell clusters suggesting they might originate and divide from a common initial cell. But this hypothesis would need to be confirmed using lineage studies. The majority of dPKD2L1 cells are found along the dorso-ventral axis close to the midline along the whole medulla-spinal axis with a homogenous density and they represent around 30% of the total PKD2L1^+^ neuron population. However, there was a differential distribution for dPKD2L1 neurons with neurons being closer to the CC and the midline in lumbSC and further away in the cervSC and Med. Most dPKD2L1 neurons are bipolar and generally extend dorsally a long dendrite (MAP2 positive) towards the CC. We observed long extensions ending with a characteristic protrusion or ‘bud’ in contact with the CC that could be followed back to dPKD2L1 cell bodies suggesting that these neurons are in contact with the CC. This hypothesis is further supported by the visualization of numerous long longitudinal PKD2L1^+^ neurites projecting along the midline to the CC that may represent dPKD2L1 neuron dendrites cut from cell bodies during section preparation. One might suggest they correspond to the population of ventral CSF-cNs’’ described by Petracca and Colleagues (2016). Nevertheless, the number of dPKD2L1 neurons showing a clear contact with the CC appears low, and one might wonder whether all observed PKD2L1^+^ cell bodies represent distal CSF-cNs. Our preliminary experiments using tracing strategies with markers (Choleratoxin B-FITC; Zhang et al., 2003) injected in the lateral ventricles support the assumption for the presence of distal CSF-cNs (data not shown) but further experiments need to be performed to specify and distinguish this cell population from the other dPKD2L1 cells and demonstrate their functional link with the CSF. We also found dPKD2L1 neurons with a more complex morphology that extend multiple dendrites in various directions but with no obvious contact with the CC and generally localized more laterally from the midline. We therefore hypothesize they would represent another subpopulation of PKD2L1^+^ neurons with either a different developmental origin or that would have migrated from the midline zone and ‘detached’ from the CC. Our present results do not provide evidence to support any of these two assumptions and further dedicated studies would be required to address this question.

Finally, using transgenic mouse (PKD-GFP and PKD-tdTomato), we provide the first evidence for their axonal projections in mice and confirm previous reports in rat and zebrafish larva. We observed thin ventral neurites running parallel and longitudinally in the rostro-caudal axis. Some of them could be seen extending from dPKD2L1 neurons. Further, in coronal sections we observed fibers bundles (either GFP^+^ or tdTomato^+^) on both side of the ventral median fissure that are MAP2 negative and strongly reminds the structure reported by Stoeckel and Colleagues (2003) in rat or by Djenoune and Colleagues (2017) in the Zebrafish larva. Therefore, we suggest they represent axons from PKD2L1^+^ neurons that extend from CSF-cNs, potentially also from dPKD2L1 neurons, to regroup ventrally along the rostro-caudal axis. As mentioned, these fiber bundles are MAP2 but also NF160 negative and we were not able to label them with the classical axonal markers (see also below) to confirm their axonal nature. This issue remains open, but crucial to be addressed in dedicated studies to confirm the axonal nature of these neurites and more importantly demonstrate the territories they innervate. This will give insight about the function of PKD2L1 expressing neurons.

In previous studies, we reported that PKD2L1 was only expressed in the soma and dendrite of CSF-cNs. Here we confirm this result on CSF-cNs and as well as for dPKD2L1 neurons but also indicate that longitudinal neurites in the ventral region are also positive for PKD2L1. Because of their localization and their negative IR against MAP2, we assume they correspond to axons and therefore PKD2L1 appear to be also present in some axons. In the experiments where we used transgenic mouse model, we also observed IR against PKD2L1 in neurites positive for GFP and tdTomato although the at a low intensity. Therefore, PKD2L1 channels appear to be expressed on axon but the functional relevance for such an expression is unknown.

### dPKD2L1 and CSF-cNs share similar phenotypical and functional properties

In mice, CSF-cNs are primarily GABAergic neurons with an intermediate maturity state observed even in 1-year-old mice (Orts-Del’Immagine et al., 2014, 2017). dPKD2L1 neurons are also GABAergic and in an intermediate maturity state (low level of NeuN along with DCX/Nkx6.1) (Orts-Del’Immagine et al., 2014, 2017; Petracca et al., 2016). Except for the PSA-NCAM IR only observed at postnatal ages, this phenotype for dPKD2L1 neurons was conserved in adult animal. Moreover, both dPKD2L1 neurons present along the midline and those localized more laterally exhibited the same phenotypical features. At the functional level, we show that dPKD2L1 cells have high input resistance, low membrane capacitance and a depolarized resting membrane potential (~-45 mV). Next, they are capable of generating APs either spontaneously or following injection of positive current pulses. Based on the discharge pattern, we find two populations with either tonic or phasic firing. In contrast, Petracca and colleagues (2016) reported that all ventral PKD2L1^+^ neurons presented a phasic firing pattern upon positive current injection. Nevertheless, their recordings were carried out on acute spinal cord slices obtained from mouse embryos (E18.5) and one can suppose, as previously reported by Marichal and colleagues (2009), that the difference observed here is due to neuronal maturation at later stages. Further, our recordings indicate that dPKD2L1 neurons receive synaptic entries with either slow or rapid decay times. Therefore, dPKD2L1 neurons appear contacted by GABAergic and glutamatergic neurons and inserted in a neuronal network that will need to be elucidated. dPKD2L1 presumed axonal processes follow a similar pathways than CSF-cNs axons (Stoeckel et al., 2003; Djenoune et al., 2017), notably the ventral midline and the rostro-caudal axis at the ventral fissure, suggesting they may have common projection path and potentially cells targets and functions. However as suggested by Djenoune and colleagues (2017) in the Zebrafish larva, ventral PKD2L1^+^ neurons might target a different pool of postsynaptic neurons and serve other function. A future challenge will be to characterize the network dPKD2L1 neurons are inserted in and identify their precise projection path and synaptic partners using targeted viral tracing strategies.

dPKD2L1 neurons exhibit spontaneous unitary current activity with properties resembling that recorded in CSF-cNs and therefore presumably carried by PKD2L1 (Orts-Del’immagine et al., 2012; Orts-Del’Immagine et al., 2016; Sternberg et al., 2018). In the present study, we did not assess the sensory properties of dPKD2L1 (pH, osmolarity or mechanic stimuli) but PKD2L1 expression might indicate they are also capable of sensing pH variation as well as other sensory stimuli. Further, our result indicates that, at least some dPKD2L1 cells, might be in contact with the CC and act as distal CSF-cNs capable of integrating signals from the CSF. On the other hand, one may suggest that the most ventral dPKD2L1 neurons, in particular those with a multipolar morphology and putatively detached from the CC, could rather act as sensors of the interstitial fluid composition to convey the collected information to yet unknown targets. This specific point would need to be addressed to assess whether these dPKD2L1 neurons are sensory neurons and what are the modalities they are sensitive to. Overall, one might wonder which function(s) dPKD2L1 serve and further study would be needed to fully address this question.

### Which cellular origin and lineage for dPKD2L1 neurons?

It has been shown that the ependymal layer forms a heterogeneous cell population and that the ventral ependymal cells represent a specific population with properties of radial glial cells and neural progenitors (Marichal et al., 2012). Further, in zebrafish (Djenoune et al., 2017) and mouse (Petracca et al., 2016; Di Bella et al., 2019), CSF-cNs were also shown to represent a heterogenous neuronal population distinguishable both phenotypically and functionally and originating from different progenitor zones. Thus, CSF-cNs in the dorso-lateral region of the CC, designated as CSF-cN’, express PKD2L1, Gata2/3, Pax6 and Nkx6.1 and are thought to originate from p2/pOL progenitor domains. In contrast, ventral CSF-cNs (designated as CSF-cN”) develop from the p3 region close to the floor plate and express PKD2L1, Gata2/3, Nkx6.1 and Nkx2.2 as well as Foxa2 (Di Bella and Colleagues, 2019). We observed that dPKD2L1 neurons, alike CSF-cN”, always contact the CC at its ventral pole, and may therefore share the same ventral origin. Moreover, early ventral CSF-cN” found around E15 are present near the SC ventral pole and might become ventral dPKD2L1 cells after fusion and/or regression of the CC ventral part. However, since only 10-20% of PKD2L1-expressing neurons are CSF-cN” around E15 (Petracca et al., 2016) and we show the number of dPKD2L1 neurons compared to CSF-cNs increases up to the early postnatal days. Therefore, we cannot exclude that new-born dPKD2L1 cells are generated at later stages from the ventral CC zone (as CSF-cNs” do) and subsequently migrate along the ventral ependymal fibers to reach their ventro-medial site. Alternatively, one can also speculate that these cells may be directly generated from neuroepithelial cells present below the CC (corresponding putatively to the remaining floor plate). Indeed, numerous PKD2L1/tdTomato positive cells are already present distally at E16 and may further migrate both ventrally and laterally after their neurogenesis. Finally, ventral CSF-cN development and maturation appears to depend on the floor plate since its absence in zebrafish mutated for cyc-1, leads to the reduction in ventral CSF-cN number and projection (Bernhardt et al., 1992). To date the developmental origin and the progenitors for PKD2L1 expressing neurons, and in particular of dPKD2L1 neurons, is not clear and further studies are still necessary. For instance, their origin and migration capabilities may be specified using organotypic cultures and/or time-lapse BrdU experiments at different developmental stage.

A recent study suggested that ventral PKD2L1^+^ cells might be the result of ectopic distribution that depends on the animal models or strains tested (Tonelli Gombalová et al., 2020). In agreement with our study, the authors reported the presence of distal PKD2L1 expressing neurons in several animal models as well as in the C57Bl6/J (the mouse line used in our study). These neurons exhibit several phenotypical characteristics largely similar to those described for dPKD2L1 neurons in our study, notably their low level or absence of NeuN expression, their location along the ventral midline, their post-natal translocation and the presence of neurons with a multipolar morphology. They further report that in C57Bl/6N mice, compared to other mice strains and rat, the proportion of distal cells relative to proximal CSF-cNs is higher and largely depends on Crumbs 1 (Crb1) and Cytoplasmic FMRP interacting protein 2 (PIR121 or Cyfip2) polymorphism observed in C57Bl/6N mice. Thus, the presence in C57Bl/6N mice of ectopic distal and largely multipolar PKD2L1^+^ neurons is partially attributed to the mutation in Crb1 and Cyfip2 genes. Nevertheless, even in C57Bl/6N, these cells were found relatively near to the CC (median value of 62 μm in the lumbar SC). In our study, we used tissue obtained from C57Bl6/J mice and observed cells much more distant in the rostral part of the CC (around 250 μm in the spMed, around 160 μm in the cMed and C1 cervSC). A similar rostro-caudal incidence was also observed for the distance to the ventral midline, that was much higher in the rostral part (Med compared to SC). Other distinctive features reported in our study for dPKD2L1 neurons include the potential link to the CC with long ventro-dorsal processes for some of the neurons, their specific astro-ependymal environment, their immature phenotype (Nkx6.1^+^, DCX^+^, PSA-NCAM^+^ in neonates), low PKD2L1 IR, their link with CSF-cNs axon bundles and their specific electrophysiological properties. Interestingly, expression of Crb1 was reported to be involved in the normal morphogenesis of CSF-cNs and to some extent the retention of their dendritic protrusion at the level of the CC (Desban et al., 2019). In the absence of native Crb1, as observed in C57Bl6/N mice, PKD2L1 expressing cells would ‘detach’ from the CC and have an altered morphology associated with an ectopic distribution. Therefore, one would suggest that, within the apical junctional complexes, expression of Crb1 is essential for CSF-cNs characteristic morphology and specific studies are necessary to specify the influence of Crb1 on distal CSF-cN morphology and distribution, their relationship with the CC and to analyze further their chemosensory functions compared to CSF-cNs.

### PKD2L1 neurons are constitutive of the ependymal stem cell niche

A common feature between dPKD2L1 neurons and CSF-cNs is their incomplete maturity state. Several studies have previously shown that CSF-cNs, despite their robust action potential discharge activity and their integration in a functional neuronal network, retain some characteristics of juvenile neurons in terms of phenotype, as shown by their expression of NKx6.1 and DCX as well as their low NeuN expression (Marichal et al., 2009; Sabourin et al., 2009; Kutna et al., 2013; Orts-Del’Immagine et al., 2014, 2017; Petracca et al., 2016). Here, we show that dPKD2L1 neurons share a similar immature state at postnatal stages that evolves with aging to an intermediate maturity status with the loss of PSA-NCAM expression like in CSF-cNs (Orts-Del’Immagine et al., 2017). Moreover, in neonates, dPKD2L1 neurons, alike CSF-cNs, have not yet acquired their definite location, as indicated by their higher proximity to the CC, suggesting that these cells may still migrate after birth (Orts-Del’Immagine et al., 2017). The fact that dPKD2L1 neurons are exclusively found in the grey matter at early postnatal ages while in adult animals they can also be found in the white matter further support this assumption. A unique feature of CSF-cNs is their close relationship with the ependymal stem cells, so that they can be considered as a constituent of the CC stem cell niche. Thus, CSF-cNs cell bodies are in the direct vicinity with ependymal cells, notably through *zona adherens* (Stoeckel et al., 2003; Hugnot and Franzen, 2011) and, using PKD2L1:GFP mouse model, we show that their axons, projecting ventro-medially, follow the ventro-medial ependymal fibers toward the ventral fissure. Here, we show that dPKD2L1 neurons are distributed along and in contact with these ependymal cell fibers and, as CSF-cNs, are present in a region enriched with GFAP positive astrocytes. Due to their astro-ependymal environment, one would therefore suggest that dPKD2L1 neurons might develop along ependymal cell fibers (see above) and be functionally connected to ependymal cells and astrocytes. It is also noticeable that CSF-cNs and ependymal cells were shown to share some molecular markers of neural progenitors such as Nkx6.1 (Fu et al., 2003), Sox2 (Petracca et al., 2016) and FoxJ1 (in zebrafish, Ribeiro et al., 2017) a result that we confirm for dPKD2L1 neurons (Nkx6.1). Thus, although the spinal stem cell niche is not capable of generating CSF-cNs in adult mammals (Marichal et al., 2009; Petracca et al., 2016), it can be suggested that this specific cell environment and potentially their contact with the CSF (Lehtinen and Walsh, 2011; Carnicero et al., 2013) may be conducive to maintain a low neuronal maturity state. Further, Di Bella and colleagues (2019) indicated that Ascl1, a basic helix-loop-helix (bHLH) transcription factor, is crucial for the differentiation of CSF-cNs. Indeed, in the absence of Ascl1, cellular development from CSF-cN progenitors is favored towards ependymal cells formation and CSF-cNs are largely absent. This result therefore suggests that CSF-Ns and ependymal cells would share common progenitors and that following SC lesion or in pathophysiological conditions Ascl1 *de nuovo* expression might allow the generation of new CSF-cNs. Further, both CSF-cNs and ependymal cells might take part in detecting lesion and inflammatory signals following trauma, neuroinflammation or neurodegenerative situation or even in regenerative processes. Nevertheless, to date, they are no evidence to support this function and dedicated studies are needed to clarify these points.

## CONCLUSION

In our study, we show that neurons expressing PKD2L1 can be found distributed along the ventro-medial axis in the medullo-spinal tissue and presumably they represent a subpopulation of CSF-cNs related to the neurons described by Petracca and colleagues (2016; CSF-cNs’’). These neurons share many phenotypical and functional properties with the more classical CSF-cNs’ and might play a role as sensory neurons either by detecting signals in the CSF or in the interstitial tissue to inform neuronal partners yet to be identified. On many aspects, CSF-cNs are a unique neuronal population and the presence of this distal subpopulation just add more complexity in the determination of their role in mammalian CNS. One of the future challenges concerning this latter population will be to determine whether they are in contact with the CSF, to identify the network they are inserted in and assess their sensory properties as well as their potential role in reparatory processes.

## ABBREVIATIONS

CC: central canal
cMed: caudal Medulla
cervSC: cervical spinal cord
CSF: cerebrospinal fluid
CSF-cN: cerebrospinal fluid-contacting neuron
DCX: doublecortin
IR: immunoreactivity
lumbSC: lumbar spinal cord
MAP-2: Microtubule-Associated Protein 2
Med: Medulla oblongata
NeuN: neuronal nuclei protein
EZ: ependymal zone
PKD2L1: polycystin kidney disease 2-like 1 protein
PSA-NCAM: polysialylated neuronal cell adhesion molecule
SC: spinal cord
spMed: sub-postremal Medulla

## ACKNOWLEDGEMENTS

This research was supported by funding obtained from Centre National pour la Recherche Scientifique (CNRS, INSB), Aix-Marseille University, the “Région Provence-Alpes-Côte d’Azur” and the “Conseil Général des Bouches-du-Rhône” (PACA, CG13 - Neuracid, JT, l’Agence National pour la Recherche et la Deutsche Forschung Gemeinshaft (ANR-DFG PRCI-MOTAC80C/A134/AN16HRJ NMF, NW). NJ was recipient of an A*Midex AMU PhD scholarship from the ‘Integrative and Clinical PhD international program’ and the ANR.

We acknowledge the Institut de Neurosciences de la Timone (INT) technical facilities for their support in the study (NeuroBioTools: Molecular Biology and Histology; INPHIM: confocal microscopy).

## Author contributions

N.J. performed electrophysiological recordings and analysis and contributed to histological studies; C.M. performed histological experiments, analyzed the data; J.T. contributed to experiment design and electrophysiological analysis; A.K. designed and performed histological experiments, analyzed the data; N.W. coordinated the study, designed and analyzed electrophysiological and histological experiments. A.K. & N.W. drafted the manuscript and all authors participated in manuscript preparation and editing.

## Data accessibility

Folders containing the measurements and statistical analysis as well as the raw images of the immunohistofluorescence experiments used for the analysis and illustrated in figures are provided. We are happy to share all other data upon request (follow link: https://cloud.int.univ-amu.fr/index.php/s/8fF9Hj96yCz9nR4).

